# The modification landscape of *P. aeruginosa* tRNAs

**DOI:** 10.1101/2024.02.21.581370

**Authors:** Mariana D. Mandler, Siddhardha S. Maligireddy, Wilfried M. Guiblet, Christina M. Fitzsimmons, Kayla S. McDonald, Delayna L. Warrell, Pedro J. Batista

**Affiliations:** Laboratory of Cell Biology, Center for Cancer Research, National Cancer Institute, National Institutes of Health; Laboratory of Molecular Biology, Center for Cancer Research, National Cancer Institute, National Institues of Health

**Keywords:** tRNA, tRNA modification, *P. aeruginosa*, 4-thiouridine, variable region, acp^3^U

## Abstract

RNA modifications have a substantial impact on tRNA function, with modifications in the anticodon loop contributing to translational fidelity and modifications in the tRNA core impacting structural stability. In bacteria, tRNA modifications are crucial for responding to stress and regulating the expression of virulence factors. Although tRNA modifications are well-characterized in a few model organisms, our knowledge of tRNA modifications in human pathogens, such as *Pseudomonas aeruginosa*, remains limited. Here we leveraged two orthogonal approaches to build a reference landscape of tRNA modifications in *E. coli*, which enabled us to identify similar modifications in *P. aeruginosa*. Our analysis revealed a substantial degree of conservation between the two organisms, while also uncovering potential sites of tRNA modification in P. aeruginosa tRNAs that are not present in *E. coli*. The mutational signature at one of these sites, position 46 of tRNA^Gln1(UUG)^ is dependent on the *P. aeruginosa* homolog of TapT, the enzyme responsible for the 3-(3-amino-3-carboxypropyl) uridine (acp^3^U) modification. Identifying which modifications are present on different tRNAs will uncover the pathways impacted by the different tRNA modifying enzymes, some of which play roles in determining virulence and pathogenicity.

## Introduction

RNA modifications have a substantial impact on gene expression, modulating the function of both coding and non-coding RNAs (Frye et al. 2018). Many RNA modifications were first identified on tRNA, which are known to have the highest frequency of modified nucleosides per RNA length, averaging eight modifications per tRNA molecule in bacteria and thirteen in eukaryotes (Pan 2018; Zhang et al. 2022). Reflecting the importance of tRNA modifications in cellular function, about 1.5% of the proteins in the *Escherichia coli* (*E. coli*) proteome are necessary for the synthesis of the different tRNA nucleotide modifications (de Crécy-Lagard et al. 2020; Björk and Hagervall 2014). On tRNA transcripts, modifications are not uniformly distributed, and the function of a given modification depends on its location. While modifications in the anticodon loop play an important role in translational fidelity, modifications in the tRNA D-arm and T-arm influence tRNA structural stability (Pan 2018).

In bacterial pathogens, tRNA modifications are important regulators of stress response and the expression of virulence factors (Koh and Sarin 2018). In *Vibrio cholerae*, the pathogen responsible for Cholera, genes required for tRNA modification, including *thiI*, while dispensable for growth under laboratory conditions, are required for optimal bacterial growth in an animal model (Pritchard et al. 2014). The enzyme ThiI is responsible for the addition of 4-thiouridine (s^4^U) at position 8 or 8 and 9, of bacteria and archaea (Dunin-Horkawicz et al. 2006; Boccaletto et al. 2022). This modification plays a unique role in prokaryote biology, as response to near-ultraviolet (UV) light is dependent on the presence of s^4^U in tRNAs (Thomas and Favre 1980; Kramer et al. 1988; Ramabhadran and Jagger 1976; Thomas et al. 1981). In addition, s^4^U has been shown to function cooperatively with other modifications to ensure that tRNAs fold properly to avoid clearance through the RNA degradosome in a tRNA quality control pathway (Kimura and Waldor 2019). In *Pseudomonas aeruginosa (P. aeruginosa)*, the tRNA modifying enzyme MiaB, is a positive regulator of the Type III secretion system and is required for induced cytotoxicity of human lung epithelial cells (Lin et al. 2022). MiaB catalyzes the final step in the synthesis of 2-methylthio-N6-isopentenyladenosine (ms^2^i^6^A), using N6-isopentenyladenosine (i^6^A) as a substrate (Esberg et al. 1999). The effect of MiaB on Type III secretion system might be independent of tRNA modification levels (Lin et al. 2022). In multiple pathogens, the tRNA uridine 5-carboxymethylaminomethyl synthesis enzyme GidA plays a critical role in the rhl-dependent quorum-sensing pathway. In a complex with MnmE, GidA catalyzes the formation of methyluridine derivates at the wobble position, critical for the decoding of NNG/A codons (Boccaletto et al. 2022; Gupta et al. 2009; Moukadiri et al. 2009). GidA homologs control expression of virulence genes, with no impact on growth, by modulating posttranscriptional regulation of essential transcriptional activators (Durand et al. 2000; Cho and Caparon 2008; Kinscherf and Willis 2002; Sha et al. 2004). It has been proposed that mRNAs with a high proportion of codons that require methyluridine derivates at the wobble position for decoding are more sensitive to changes in expression of the GidA enzyme (Lin et al. 2022). In *P. aeruginosa*, the response to oxidative stress is also dependent on tRNA modifications. Loss of the protein TtcA, responsible for biosynthesis of 2-thiocytidine (s^2^C), tRNA methyltransferase TrmJ, responsible for 2’-O-methylcytidine (Cm), 2’-O-methyluridine (Um) and 2’-O-methyladenosine (Am), or TrmB, responsible for 7-methylguanosine (m^7^G) modification at position 46 results in hypersensitivity to oxidative stress (Romsang et al. 2018; Jaroensuk et al. 2016; Thongdee et al. 2019). The effect of TtcA, which is upregulated in response to oxidative stress, on gene expression occurs at the post-transcriptional level and loss of this enzyme impacts virulence on a host model (Romsang et al. 2018). Loss of TrmB impacts translation of Phe- and Asp-enriched mRNAs, including catalases that breakdown hydrogen peroxide, KatA and KatB. These examples highlight the role for tRNA modifications in the virulence of pathogenic bacteria.

*P. aeruginosa, a* gram-negative bacterium, is a common environmental microorganism known to, under specific circumstances, become an opportunistic pathogen capable of infecting immunocompromised individuals (Lyczak et al. 2000). This microorganism, frequently found in multi-species infections, leverages multiple virulence strategies and infects a wide range of hosts (Lyczak et al. 2000; Khare and Tavazoie 2015; Stover et al. 2000). Previous studies have explored, at a global level, RNA modifications in *P. aeruginosa* tRNAs by LC-MS/MS (Jaroensuk et al. 2016; Grobe et al. 2019) and identified modifications that are conserved between *E. coli* and *P. aeruginosa*, as well as modifications that are present exclusively in *E. coli* or *P. aeruginosa* (Jaroensuk et al. 2016; Grobe et al. 2019).

In this study, we aimed to map tRNA modifications in *P. aeruginosa* at nucleotide resolution, and determine which modification are present at each tRNA transcript. To achieve this, we employed a combination of cDNA misincorporation and termination sequencing analysis to identify sites of tRNA modification with previously established methanethiosulfonate (MTS) chemistry to enrich for s^4^U-modified RNA (Supplemental Figure 1A). Using *E. coli* as a reference, we determined which tRNA modifications are conserved between the two organisms and assigned modification status to multiple tRNA positions in *P. aeruginosa*. In addition, we identified modification sites that are present in *P. aeruginosa* but not in *E. coli*, such as position 46 of tRNA^Gln1^ and position U20B of tRNA^Glu1^, positions recently described to be modified with an acetylated form of *N*3-(3-amino-3-carboxypropyl)-uridine (acp^3^U) in *Vibrio cholerae*. Finally, we identified the *P. aeruginosa* homolog of TapT and demonstrated that the modification signature at position 46 of tRNA^Gln1^ is dependent on the activity of this enzyme. It remains to be determined if acp^3^U at this position is further modified to acetylated acp3U (acacp3U). Altogether, the approach presented here offers the discovery of sites of tRNA modifications from understudied bacteria that are implicated in their virulence and pathogenesis.

**Fig. 1.**
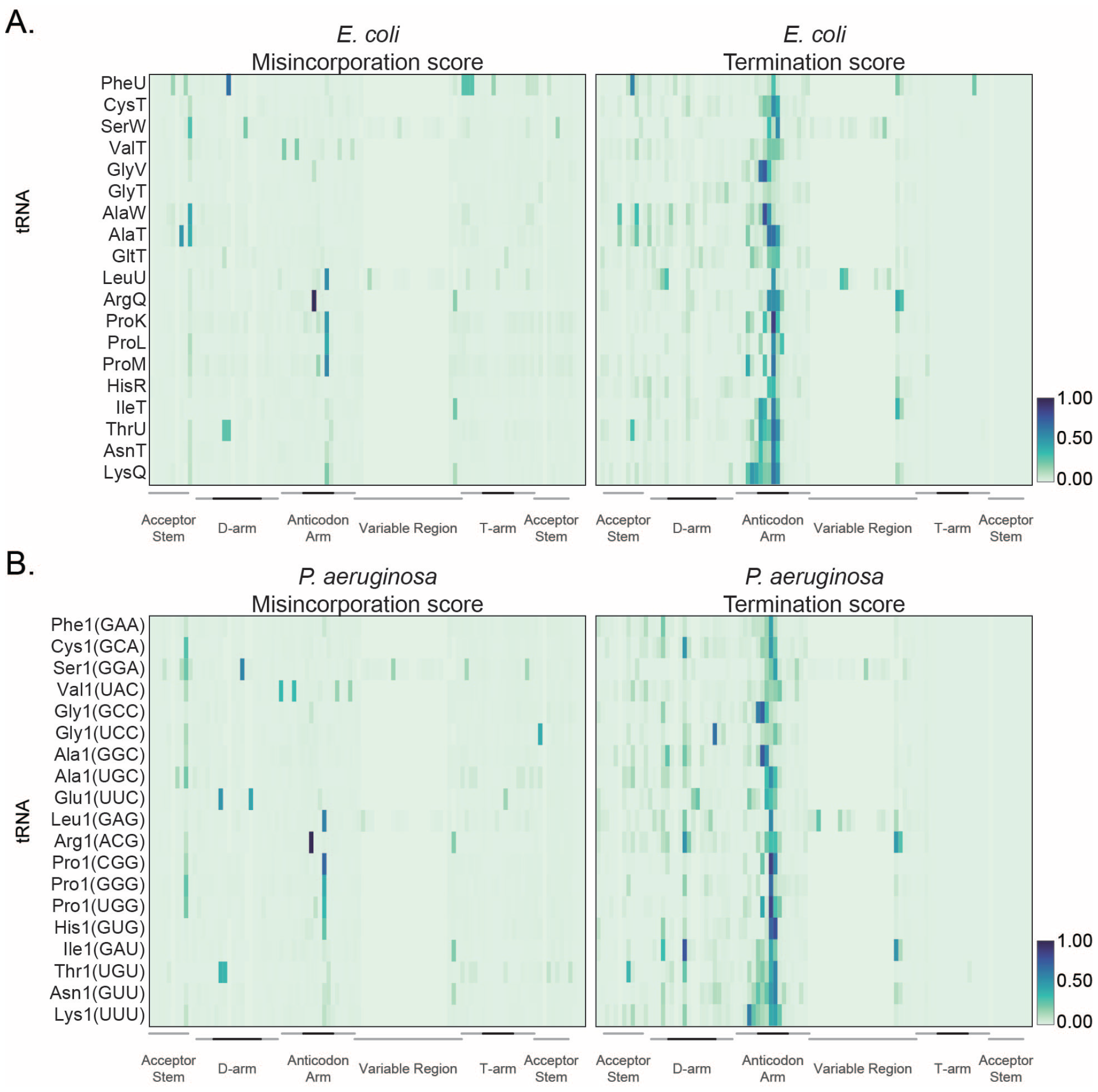
An *E. coli* reference landscape for tRNA modifications. **(A)** Heatmap of average frequency (from three independent samples) of misincorporation (left) or termination score (right) at each position for *E. coli* tRNAs extracted from cells cultured in chemically modified media and collected at (OD)_600_ =0.6. Only tRNAs with read coverage > 100 at each position and with an identified isoacceptor homolog in *P. aeruginosa* are represented. The color key at the right represents the frequency of misincorporation or termination score. Different regions of the tRNA are represented at the bottom from the 5^*′*^ (left) to the 3^*′*^ end (right) of the tRNA. Darker lines indicate the loop in that region. **(B)** Heatmap of average frequency (from three independent samples) of misincorporation (left) or termination score (right) at each position for *P. aeruginosa* tRNAs extracted from cells cultured in chemically modified media and collected at (OD)_600_ =0.6. Only tRNAs with read coverage > 100 at each position and with an identified isoacceptor homolog in *E. coli* are represented. The color key at the right represents the frequency of misincorporation or termination score. Different regions of the tRNA are represented at the bottom from the 5^*′*^ (left) to the 3^*′*^ end (right) of the tRNA. Darker lines indicate the loop in that region. The GtRNAdb names are used for *P. aeruginosa* tRNAs. The anticodon is shown in parentheses.

## Results

To identify RNA modifications in *P. aeruginosa* tRNAs, we first built a reference modification landscape in the well-characterized model organism *E. coli*. To this end, we leveraged two orthogonal approaches: analysis of cDNA mutational signature and, targeted enrichment of s^4^U modified RNAs. During cDNA synthesis, the presence of modifications that interfere with the activity of the reverse transcriptase (RT) enzyme can result in the incorporation of mismatched bases in the cDNA or premature termination of cDNA synthesis. By examining these occurrences at each position on the tRNA, defined here as mutational signature, we can effectively predict the presence of modifications (Schwartz et al. 2018; Cozen et al. 2015; Zheng et al. 2015; Richter et al. 2018; Vandivier et al. 2019; Kimura et al. 2020; Wang et al. 2021; Gogakos et al. 2017). Due to the high frequency of complex modifications found in the anticodon loop of tRNAs transcripts, cDNA products are often truncated. Consequently, read coverage for positions in proximity to the 5^*′*^ end, where s^4^U is found (Bommisetti and Bandarian 2022), may be restricted. To address this limitation, we employed a strategy to isolate tRNAs through affinity purification targeting the s^4^U modification to determine levels of s^4^U modification (Duffy et al. 2015).

### Generation of a tRNA reference map in E. coli

To generate the reference landscape of tRNA modifications in *E. coli* we analyzed tRNAs extracted from cells cultured in Luria-Bertani (LB) media or chemically defined media (CDM) and collected during exponential growth phase (optical density (OD)_600_=0.6). In order to identify both misincorporation and early termination events during cDNA synthesis, we adapted the library generation protocol described by (Flynn et al. 2016). tRNAs were gel purified from a urea polyacrylamide gel (Urea-PAGE) after ligation of an oligo adaptor to the 3^*′*^ end of total RNA. cDNA was generated with thermostable group II intron reverse transcriptase (TGIRT) and an oligonucleotide including universal molecular identifiers (UMIs), which allows for the accurate removal of PCR duplicate reads (Zarnegar et al. 2016). To ensure that truncated cDNA products are represented in the final library, the cDNA product was circularized, and the final amplicon for sequencing was generated by PCR. Reads were mapped to a custom curated transcriptome, which includes annotated coding and non-coding RNAs. In this curated transcriptome, tRNAs that share 100 percent sequence identity were grouped together (Supplemental Table 1). Reads were mapped with Bowtie 2 (Langmead and Salzberg 2012) with the inclusion of the –local parameter to account for the addition of non-templated nucleotides by TGIRT at the 3^*′*^ end of the cDNA (Qin et al. 2016). We limited our analysis to single mapped reads that include the CCA sequence at the 3^*′*^ end. The frequency of misincorporation and a termination score were calculated for each tRNA position, with the termination score determined using the formula:

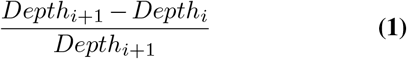

as previously described (Wang et al. 2021). Each analysis was performed on three biological replicates that show high degree of correlation (Supplemental Figure 1B). Consistent with previous studies (Schwartz et al. 2018; Cozen et al. 2015; Zheng et al. 2015; Richter et al. 2018; Vandivier et al. 2019; Kimura et al. 2020; Wang et al. 2021; Gogakos et al. 2017), we observe misincorporation, early termination, or a combination thereof (mutational signature) across all tRNAs in both culture media (Figure 1A, Supplemental Figure 2). In this study, we defined a tRNA position as modified if it met the criteria of having a read coverage equal to or greater than 100 reads and exhibits a minimum of 5 percent misincorporation or termination. Applying these criteria, we identified a mutational signature for known modifications occurring on position 8 (Figure 1A, Supplemental Figure 2, Supplemental Table 2, Supplemental Figure 4A), the anticodon loop (Supplemental Figure 3, Supplemental Figure 5, Supplemental Table 4) and the variable loop (Supplemental Figure 3, Supplemental Figure 6, Supplemental Table 5). Additionally, we detect regions of misincorporation, early termination, or both in regions without known modifications, including the anti-codon loop of certain tRNAs (Supplemental Table 2). It is worth noting that with our assay conditions, there are several types of modifications for which a consistent mutational signature is not observed (Supplemental Figure 3, Supplemental Table 2). We observe that misincorporation at position 8 is dependent upon the enzyme ThiI, confirming that the detected mutational signature is dependent upon the presence of s^4^U modification (Supplemental Figure 4A). Read coverage at position 8 is often low due to early termination of cDNA synthesis at the anticodon region. Therefore, we employed MTSEA capture as an orthogonal method to characterize the s^4^U modification status of *E. coli* tRNAs. As previously demonstrated, MTSEA reacts with RNA extracted from wild type *E. coli*, but not with RNA extracted from cells lacking the enzyme tRNA sulfurtransferase (ThiI) (Duffy et al. 2015; Mueller et al. 1998) (Supplemental Figure 4B, 4C). MT-SEA enriched RNAs isolated from wild type *E. coli* (strain BW25113) migrated at the size range that corresponds to tR-NAs (Favre et al. 1971; Emilsson et al. 1992) (Supplemental Figure 4C). Among the tRNAs detected in this experiment, 13 and 11 were significantly enriched (log2foldchange (l2fc) > 0.6 and padj < 0.05) in RNA collect from cells cultured in LB and chemically defined media, respectively (Supplemental Figure 4D, Supplemental Table 3A and 3B). The MTSEA enrichment scores accurately reflected the misincorporation analysis, wherein frequency of misincorporation at position 8 correlated with MTSEA-dependent enrichment (Pearson correlation coefficient R = 0.53, and 0.55) (Supplemental Figure 4E). These results suggest that both MTSEA capture and mu-tational analysis can be used to identify sites of s4U modification in tRNAs. The combination of targeted capture of s4U and analysis of mutational signature in the well-characterized model organism *E. coli* allowed us to build a reference map to assign RNA modifications to *P. aeruginosa* tRNAs.

**Fig. 2.**
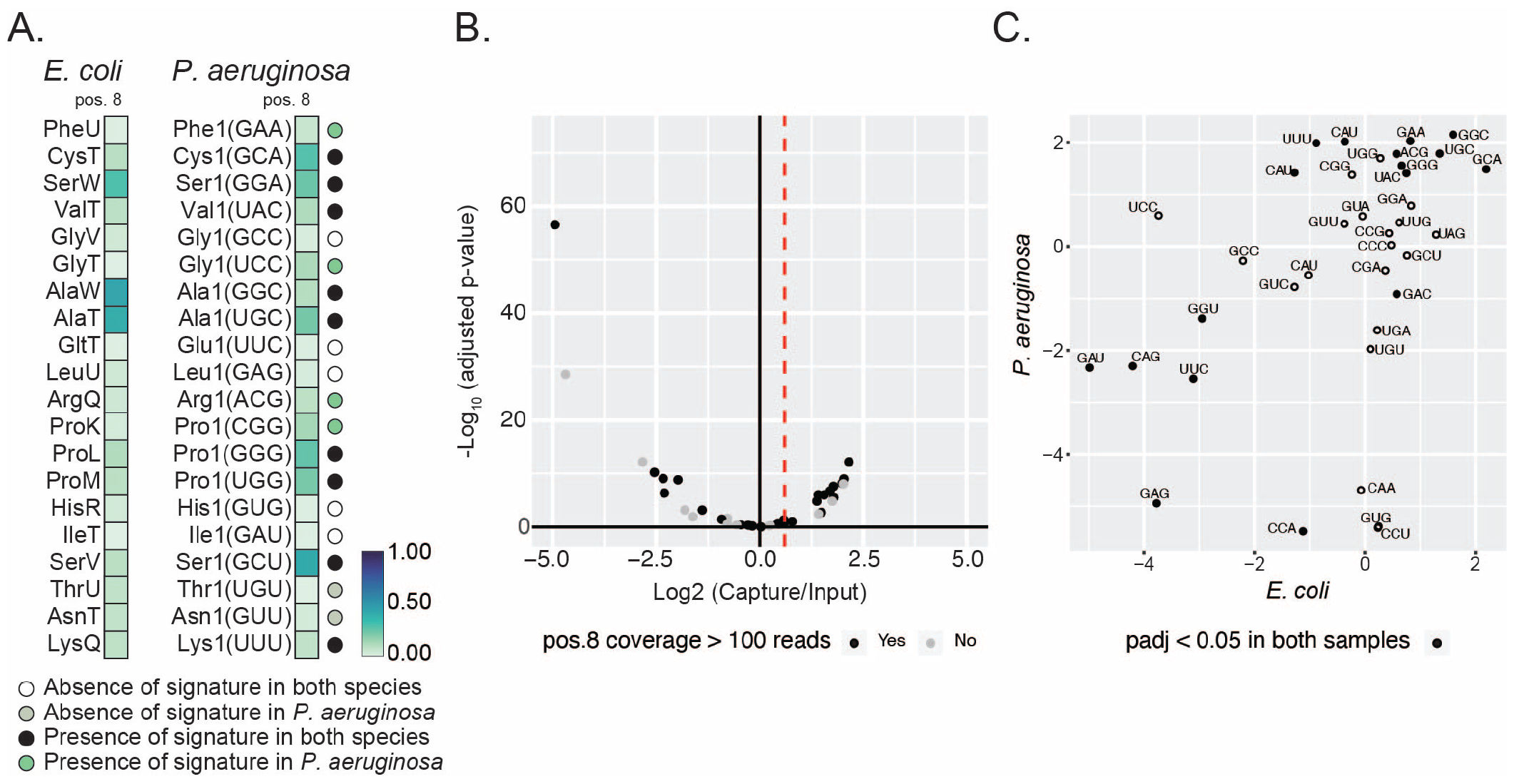
Conservation of s^4^U between *E. coli* and *P. aeruginosa*. **(A)** Heatmap of average frequency (from three independent samples cultured in chemically defined media) of misincorporation at position 8 of isoacceptor homologue tRNAs in *E. coli* and *P. aeruginosa*. The color key at the right represents the frequency of misincorporation. Only tRNAs with read coverage > 100 at position 8 and with an identified isoacceptor homolog in both species are represented. The absence or presence of a signature in both species or *P. aeruginosa* based on a five percent cutoff of misincorporation is colored coded in circle marks by the key listed below the map, and labeled on the right of the map. The criteria are based on the nature of the annotated modification. The GtRNAdb names are used for *P. aeruginosa* tRNAs. The anticodon is shown in parentheses. **(B)** Volcano plot representation of log 2-fold change in abundance after MTSEA capture of tRNA from *P. aeruginosa* cells. Dotted red line indicates the threshold used to define a tRNA as enriched (l2fc > 0.6). Color indicates if read coverage at position 8 is above 100 in misincorporation analysis. **(C)** Scatter plot representation of log 2-fold change in abundance after MTSEA capture for isoacceptor homologue tRNAs from *E. coli* and *P. aeruginosa*. Full circle indicates tRNAs that are significantly enriched in both samples.

**Fig. 3.**
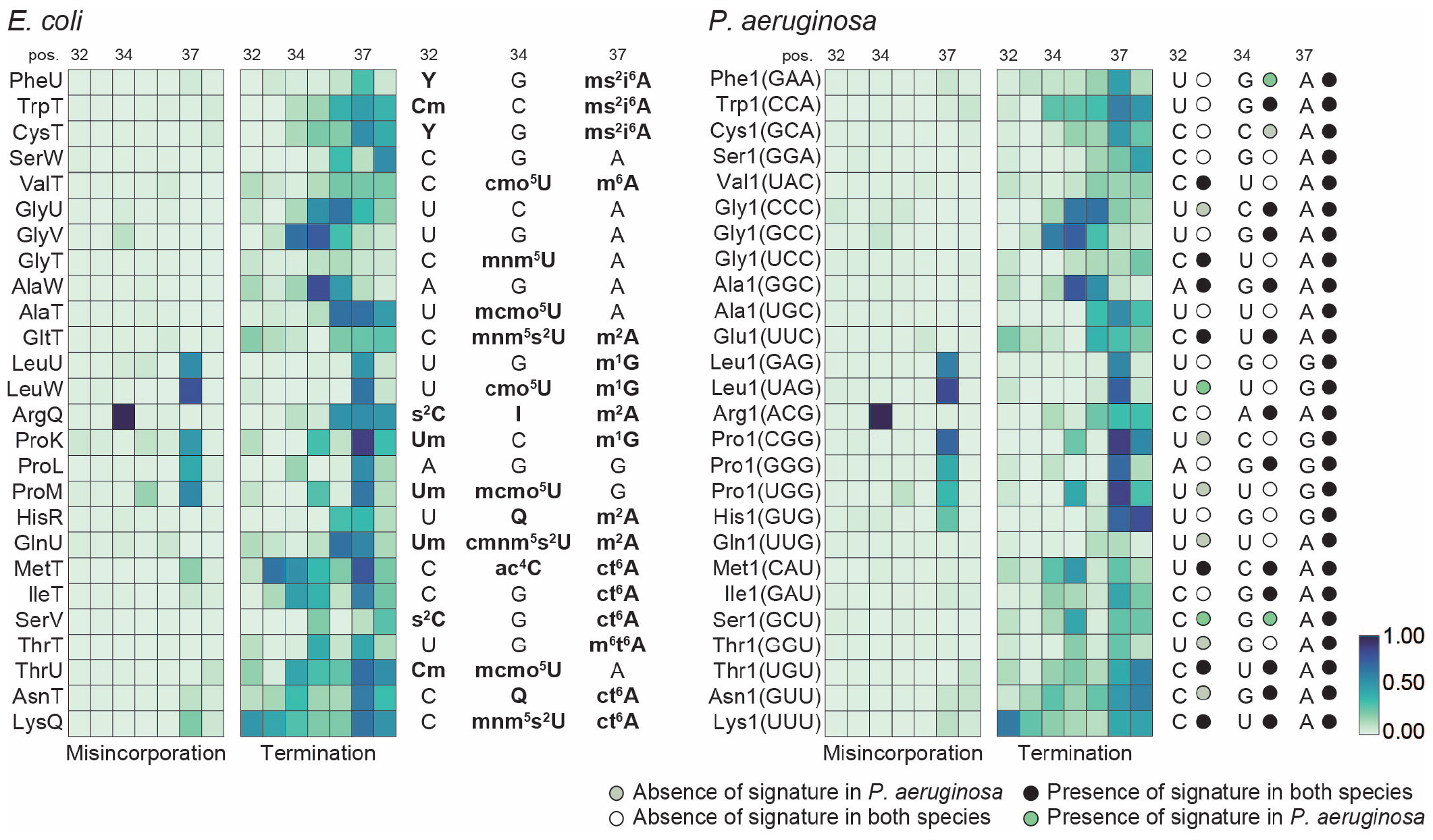
Conservation of tRNA modifications at the anticodon region. **(A)** Heatmap representation of average frequency of misincorporation and average termination score between positions 32 and 38 in tRNA from *E. coli* (left) and *P. aeruginosa* (right). Only tRNAs with read coverage > 100 in the anticodon loop and with an identified isoacceptor homolog in the two species are represented. Positions 32, 34 and 37, known to be modified are annotated at the top of each heatmap. Annotated modifications for *E. coli* are represented to the right of the *E. coli* heatmap. The short name for each modification is used (Supplemental Table 11). Sequences in *P. aeruginosa* at positions 32, 34 and 37 are represented to the right of the *P. aeruginosa* heatmap. The GtRNAdb names are used for *P. aeruginosa* tRNAs. The anticodon is shown in parentheses. The absence or presence of a signature in both species or *P. aeruginosa* is based on a five percent cutoff of either misincorporation or termination for positions 32, 34 or 37. Circles are colored coded by the key listed below the map and labeled on the right of the map. The criteria is based on the nature of the annotated modifications. The color key at the bottom right represents the frequency of misincorporation or termination score.

**Fig. 4.**
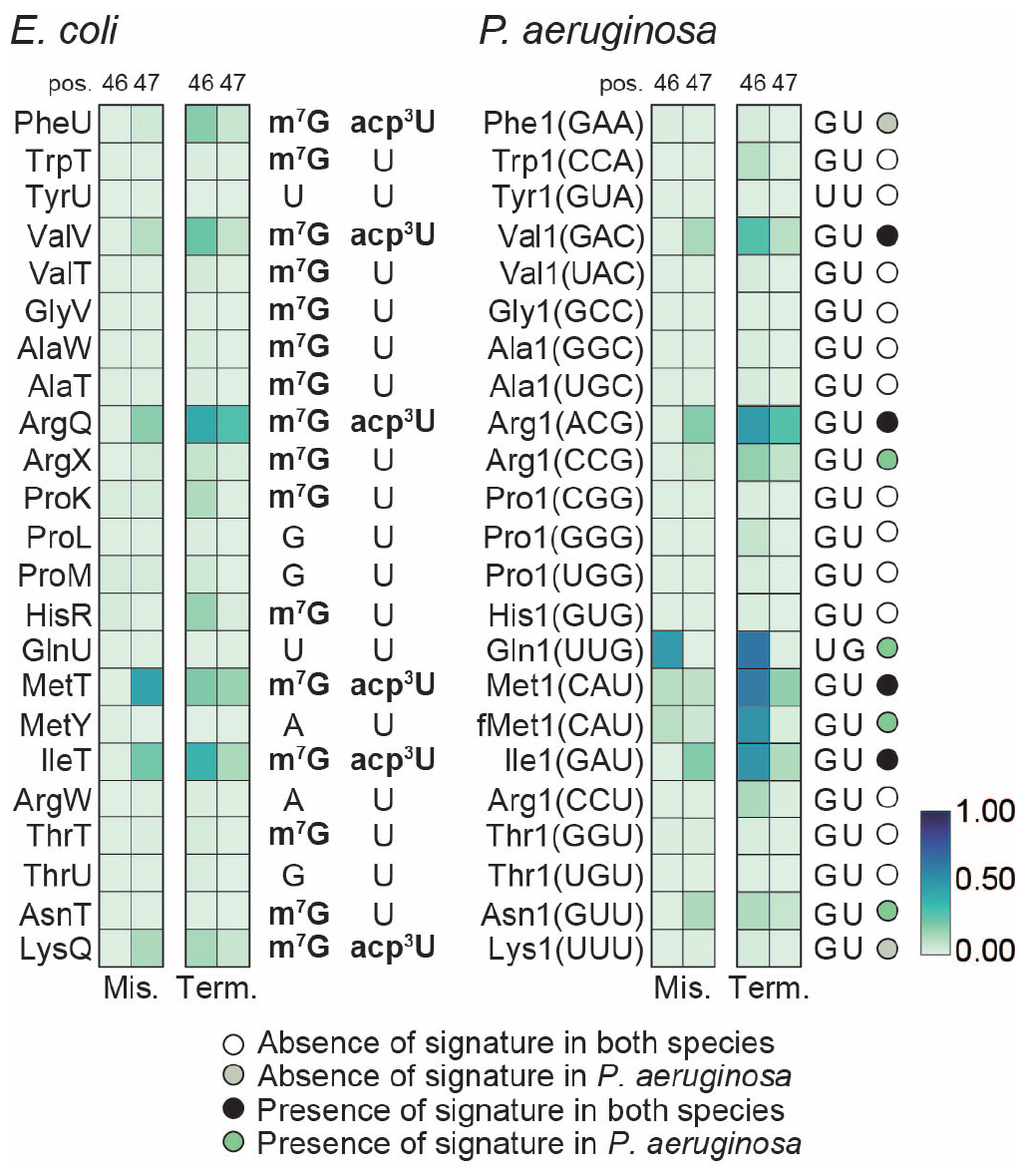
Conservation of acp^3^U and m^7^G *E. coli* and *P. aeruginosa*. **(A)** Heatmap representation of average frequency of misincorporation and average termination score at positions 46 and 47 in tRNA from *E. coli* (left) and *P. aeruginosa* (right). Only tRNAs with read coverage > 100 at position 46 and with an identified isoacceptor homolog are represented. Known modifications for *E. coli* are represented to the right of the *E. coli* heatmap. The short name for each modification is used (Supplemental Table 11). Sequences in *P. aeruginosa* at positions 46 and 47 are represented to the right of the *P. aeruginosa* heatmap. The absence or presence of a signature in both species or *P. aeruginosa* based on a five percent cutoff of both misincorporation and termination for positions 46 or 47 is colored coded in circle marks by the key listed below the map, and labeled on the right of the map. The criteria is based on the nature of the annotated modifications. The color key at the bottom right represents the frequency of misincorporation or termination score.

**Fig. 5.**
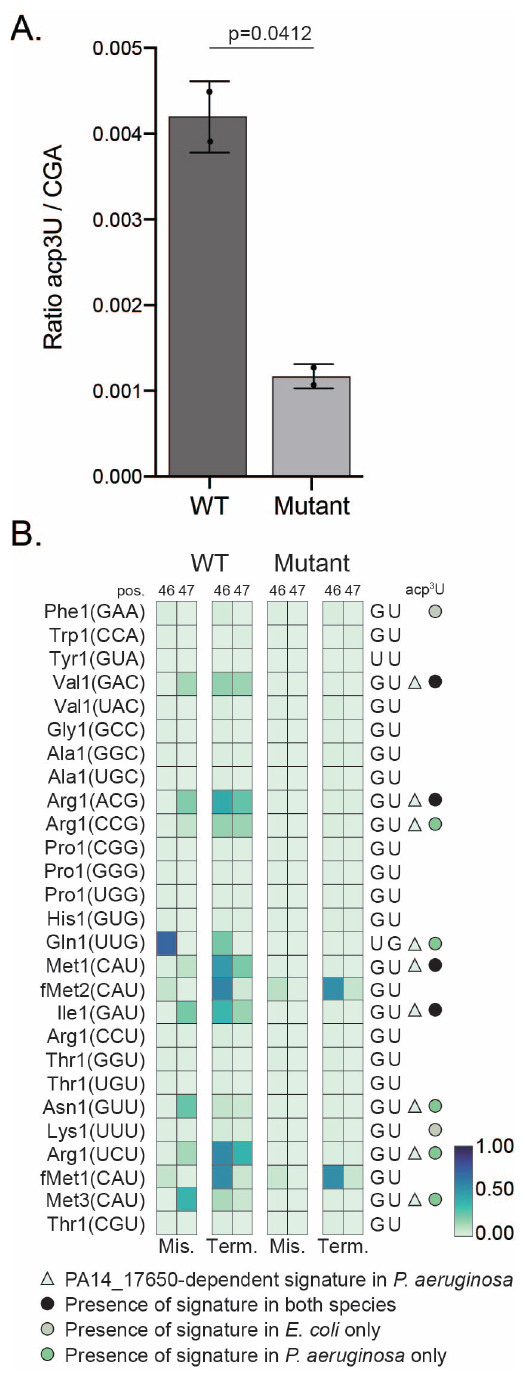
Modification sites at positions 46 or 47 that are lost in mutant strain of *P. aeruginosa* PA14_17650. **(A)** LC-MS/MS quantification of acp^3^U from purified tRNA of wild-type *P. aeruginosa* versus PA14_17650 mutant for *n*=2 replicates. **(B)** Heatmap representation of average frequency of misincorporation and average termination score at positions 46 and 47 in tRNA of *P. aeruginosa* from wild-type (left) and PA14_17650 mutant (right). Only tRNAs with read coverage > 100 at position 46 are represented. The GtRNAdb names for *P. aeruginosa* tRNAs are listed on the left, with the anticodon shown in parentheses. The short name for the modification is shown at top right column (Supplemental Table 11). The triangle symbol labeled on the right of the map indicates whether a signature is dependent PA14_17650, wherein the signature is lost in the mutant. The circle marks labeled on the right of the map are color coded by a conserved signature, presence only in *E. coli*, or novel signature that is present in *P. aeruginosa* using the key listed below the map. The criteria used is based on a five percent cutoff of both misincorporation and termination for either positions 46 or 47. The color key at the bottom right represents the frequency of misincorporation or termination score.

### Establishing a tRNA modification map for P. aeruginosa

Previous studies have cataloged RNA modifications present in *P. aeruginosa* tRNAs by mass spectrometry (Jaroensuk et al. 2016; Grobe et al. 2019) and identified modifications that are conserved between *P. aeruginosa* and *E. coli*, as well as modifications that are found exclusively in each species. We aimed to assign globally conserved modifications between *P. aeruginosa* and *E. coli* using our data. For this comparison, tRNA transcripts were matched on the basis of both anticodon and sequence similarity (Supplemental Table 6). In order to assign conserved modifications to specific tRNA positions, we compared the pattern of misincorporation, early termination or both observed in tRNA sequencing libraries of both species (Figure 1A, 1B, Supplemental Figure 7, Supplemental Table 7). While the patterns of misincorporation and termination are generally similar, we identified some notable differences.

### s^4^U modification is present in P. aeruginosa tRNAs

In *P. aeruginosa* we detect a mutational signature at position 8, here defined as misincorporation of at least 5% (Figure 1B, Figure 2A), similar to what we observed in *E. coli*, which suggests conservation of s^4^U between the two species. However, we noted several exceptions: tRNA^ThrU(UGU)^ and tRNA^AsnT(GUU)^ only have a signature in *E. coli*, while tRNA^Phe1(GAA)^, tRNA^Gly1(UCC)^, tRNA^Arg1(ACG)^, and tRNA^Pro1(CGG)^ only have a signature in *P. aeruginosa*. As described for our *E. coli* experiments, coverage at this position in our datasets is under 100 reads for several tRNAs. Using MTSEA capture we identified 13 significantly enriched tRNAs (l2fc > 0.6 and padj < 0.05) (Figure 2B, Supplemental Table 8). The mutational signature at position 8 of P. aeruginosa tRNAs correlates positively with enrichment after MTSEA capture (Pearson correlation coefficient R = 0.65) (Supplemental Figure 8, Supplemental Table 9). Overall, we observe a similar pattern of s^4^U enrichment between *E. coli* and *P. aeruginosa* tRNAs (Figure 2C, Supplemental Table 10), supporting the idea that this modification is conserved between the two species.

### Similarities and differences of tRNA modification between E. coli and P. aeruginosa in the anticodon region

When comparing misincorporation and early termination in the anticodon regions of *E. coli* and *P. aeruginosa* we note a significant conservation of mutational signatures for the majority of modifications. Importantly, all signatures at position 37 are conserved in both species, which includes the modifications for 2-methylthio-N-6-isopentenyl adenosine (ms^2^i^6^A), 2-methyladenosine (m^2^A), 1-methylguanosine (m^1^G), cyclic N6-threonylcarbamoyladenosine (ct^6^A) and N6-methyl-N6-threonylcarbamoyladenosine (m^6^t^6^A) (Figure 3). Interestingly, we find tRNAs in *P. aeruginosa* with a mutational signature that is conserved in the *E. coli* homologous tRNAs, for which no modifications have been annotated, such as for tRNA^Gly(CCC)^, tRNA^Gly(GCC)^ and tRNA^Ser(GGA)^ (Figure 3), which highlights the need to continue to annotate tRNA modifications, even in well characterized organisms. As with our analysis of the s^4^U modification at position 8, we observe several exceptions. For example, *E. coli* tRNA^GlnU(UUG)^ has a much stronger signature than the one observed in *P. aeruginosa* for the homologous tRNA, suggesting the presence of a different type of modification at position 37. In some cases, a drop in coverage at position 37 due to modifications that cause strong termination limits us from determining how well modifications are conserved at upstream positions, such as position 32. We also found examples where a mutational signature is observed in both species at the same position, even though the nucleotide sequence is different: m^2^A at position 37 of tRNA^HisR(GUG)^ and m^1^G at position 37 of tRNA^ArgX(CCG)^ in *E. coli* correspond to G and A in the *P. aeruginosa* homologous tRNAs. Finally, we also see conserved signatures at nucleotides which have different primary sequences and are not annotated as modified, such as position 32 of tRNA^MetT(CAU)^ and tRNA^Met1(CAU)^ (Supplemental table 7). This observation implies that different modifications are present at these positions.

### Similarities and differences in tRNA variable region between E. coli and P. aeruginosa

The variable region of *E. coli* tRNAs features two well-defined modifications at positions 46 and 47, namely m^7^G and acp^3^U. In *E. coli* we detected a mutational signature for all sites annotated to be modified with both m^7^G and acp^3^U (Figure 4). However, this mutational signature is absent in the majority of tRNAs annotated to be solely m^7^G modified, suggesting that the observed mutational signature is derived from the presence of acp^3^U. This is in line with previous observations that m^7^G alone lacks a mutational signature (Wang et al. 2021). In homologous *P. aeruginosa* tRNAs, we do not observe a mutational signature, suggesting that the absence of acp^3^U is conserved. The exceptions are tRNA^Arg1(CCG)^, tRNA^Asn1(GUU)^ and tRNA^Arg1(UCU)^ where presence of a mutational signature suggests the presence of the acp^3^U modification in *P. aeruginosa* (Supplemental Table 7, Figure 4, and Figure 5). Notably, the tRNA^Arg1(UCU)^ nucleotide sequence differs from tRNA^ArgU(UCU)^ at position 47, where the U required for the acp^3^U modification is only present in *P. aeruginosa*. For tRNAs annotated to carry both m7G and acp3U in *E. coli*, we observe a conserved mutational signature for tRNA^Val1(GAC)^, tRNA^Arg1(ACG)^, tRNA^Met1(CAU)^ and tRNA^Ile1(GAU)^, suggesting acp^3^U is present in the *P. aeruginosa* homolog tRNAs. Absence of a mutational signature for tRNA^Phe1(GAA)^ and tRNA^Lys1(UUU)^ in *P. aeruginosa* implies that these tRNAs are not acp^3^U modified (Figure 4). We observed a mutational signature for *P. aeruginosa* tRNA^Gln1(UUG)^ and tRNA^fMet1(CAU)^ which are not annotated to be modified in *E. coli*. In both cases, the primary sequence is different between the two species (Figure 4).

### The Variable region mutational signature is dependent on PA14_17650 (tapT)

To confirm that the mutational signature we observe at positions 46 and 47 in *P. aeruginosa* derives from the presence of acp^3^U, we aimed to map tRNA modifications in a mutant lacking the enzymatic activity required for this modification. In *E. coli*, acp^3^U is deposited by TapT, an enzyme containing a DTW domain (Meyer et al. 2020; Takakura et al. 2019). The *P. aeruginosa* protein with highest level of homology to *E. coli* TapT is PA14_17650, the ortholog of PAO-1 YfiP (Takakura et al. 2019) (Supplemental Table 12). We obtained a mutant for PA14_17650 from a nonredundant library of *P. aeruginosa* strain PA14 transposon insertion mutants (Liberati et al. 2006). Liquid Chromatography tandem mass spectrometry (LC-MS/MS) analysis of tRNA isolated from a PA14_17650 mutant strain confirms a role in acp^3^U deposition for this enzyme (Figure 5A). Before generating sequencing libraries for the PA14_17650 mutant strain and a matched wild type control, we tested multiple reverse transcriptase enzymes for sensitivity to modifications at this position. TGIRT generated the strongest signature at a site predicted to be acp^3^U modified (Supplemental Figure 9). As observed in previous experiments, the correlation between mutational signatures across replicates was high for the tRNA sequencing libraries generated from the PA14_17650 mutant strain and the matched wild type (Supplemental Figure 10). Comparison between wild type *P. aeruginosa* and the PA14_17650 mutant reveals a loss of misincorporation and termination signal specifically at position 46 and 47 (Supplemental Figure 11 and 12, Supplemental Table 13). This observation supports the idea that the m^7^G does not contribute to this mutational signature and establishes PA14_17650 as the *P. aeruginosa* homolog of *E. Coli* TapT. The mutational signature on most tRNAs predicted to be acp^3^U modified is lost in the absence of PA14_17650, confirming these tRNAs carry either the acp^3^U modification or a derivate, in *P. aeruginosa*. The two exceptions are tRNA^fMet1(CAU)^ and tRNA^fMet2(CAU)^, suggesting that modifications at these positions are deposited by a different enzyme. Interestingly, the mutational signature observed at position 46 of tRNA^Gln1(UUG)^ was also lost in the PA14_17650 mutant, showing that modification at this position requires PA14_17650 activity. This position was previously found to be acacp^3^U modified in *Vibrio cholerae*, where the primary tRNA sequence is these positions is UG, the same as *P. aeruginosa* (Kimura et al. 2020). However, we were not able to identify a clear homolog of AcpA from *V. cholerae* in *P. aeruginosa* (Supplemental Table 12). In *Vibrio cholerae*, this modification is also present at position 20 of tRNA^Glu1(UUC)^. In *P. aeruginosa* we also observe a mutational signature at this position, but this signature is not dependent on PA14_17650 activity. This position could be acp^3^U modified by a different enzyme or have an unrelated modification. Further work is necessary to determine what is the nature of the modification present at position 20 of tRNA^Glu1(UUC)^, position 47 of tRNA^fMet1(CAU)^ and tRNA^fMet2(CAU)^, and whether acp^3^U that is found at position 46 in tRNA^Gln1(UUG)^ can be subsequently modified to acacp^3^U.

## Discussion

RNA modifications expand the functional repertoire of each nucleoside, having a profound impact on RNA function. On tRNAs, RNA modifications play important roles in stabilizing secondary and tertiary structure and ensuring fidelity during translation (reviewed in (de Crécy-Lagard and Jaroch 2021)). Across prokaryotic pathogens, tRNA modifications play critical roles in regulating stress responses, adaptation to the host environment and expression of virulence factors (Koh and Sarin 2018). The repertoire of tRNA modifications in *Pseudomonas aeruginosa*, an opportunistic pathogen, has been characterized using mass spectrometry (Jaroensuk et al. 2016; Grobe et al. 2019). However, a comprehensive understanding of the specific tRNA modifications present in each tRNA molecule is currently lacking. Determining which modifications are present on different tRNAs can help elucidate the role of enzymes involved in tRNA modification on bacterial survival and pathogenicity (Duechler et al. 2016). Here, we leveraged the well-characterized *E. coli* tRNA modification landscape to predict tRNA modifications in *P. aeruginosa*. Furthermore, we identified PA14_17650 as the gene responsible for acp^3^U deposition in the variable loop of *P. aeruginosa* tRNAs.

### A reference map in E. coli

In this study, we combined chemical capture and cDNA misincorporation analysis to build a reference map of tRNA modifications in *E. coli* with the objective of identifying modifications that are conserved in *P. aeruginosa* tRNAs. While analysis of RT misincorporation involves less sample manipulation, and provides information on a wide range of modifications, the chemical capture approach provides higher sensitivity for modifications of interest, overcoming limitations such as low read coverage at the modification site, lack of a mutational signature for a given modification, or low mutational rate. We find that the two approaches can be used as orthogonal validation for modifications of interest. Other approaches such as mass spectrometry or the use of mutant strains lacking the enzymatic activity for the modification in question can be used to validate findings from sequencing-based approaches, as we demonstrate in this study for acp^3^U. In *E. coli*, mutational signatures occur predominantly at the anticodon loop, and most positions with a mutational signature overlap with an annotated tRNA modification. We also detected mutational signatures in regions where a modification has not yet been annotated and might infer novel sites of modification worth further exploration. Once we generated a reference landscape of tRNA modifications in *E. coli*, we analyzed our *P. aeruginosa* tRNA sequencing data sets generated in parallel, to determine which modifications are conserved between the two species. Our data supports conservation of several modifications between *E. coli* and *P. aeruginosa*.

### Anticodon loop modifications in P. aeruginosa

We identified several modifications in the anticodon loop to be conserved between *E. coli* and *P. aeruginosa* (Figure 3). Some of these modifications are synthesized by en-zymes linked to virulence phenotypes while others are deposited by enzymes required for survival. One example is ms^2^i^6^A at position 37. Synthesis of ms^2^i^6^A is a multistep reaction that requires the activity of MiaA, which converts A to isopentenyl adenosine (i^6^A), and MiaB which converts i^6^A to ms^2^i^6^A. While previous mass spectrometry studies did not detect this modification in *P. aeruginosa* (Jaroensuk et al. 2016; Grobe et al. 2019), Lin *et al*. (Lin et al. 2022) demonstrated that loss of MiaB in the PAO1 strain leads to accumulation of i^6^A and a slight decrease in ms^2^i^6^A, suggesting the presence of this modification and potential alternative pathways for modifying i^6^A in *P. aeruginosa*. Alternately, ms^2^i^6^A might be further modified in *P. aeruginosa*. A protein with 39.92% sequence identity to *S. typhimurium* MiaE, which converts ms^2^i^6^A to ms^2^io^6^A (Persson and Björk 1993), can be found in the *P. aeruginosa* genome (Supplemental Table 12) (Jaroensuk et al. 2016) and might play a role in this pathway. Additionally, we observed strong conservation of the modified inosine at position 34 of tRNA^ArgQ(ACG)^, which is synthesized by TadA, an essential gene in *E. coli* (Wolf et al. 2002). Other modifications known to play important roles in translation fidelity and protein homeostasis such as ct^6^A, deposited by TsaD (Su et al. 2022), and m^6^t^6^A, deposited by TrmO (Kimura et al. 2014) are also conserved. The absence of the methyl group of m^6^t^6^A37 in tRNA^ThrV(GGU)^ reduces the efficiency of this tRNA to read its cognate codon (Qian et al. 1998). Several modifications at the anticodon, such as the *P. aerug-inosa*-specific 2^*′*^-O-methyladenosine (Am) modification at position 32 of tRNA^Pro1(GGG)^ (Jaroensuk et al. 2016), do not generate a clear mutational signature. For these modifications we were not able to determine conservation between the two organisms, which is a limitaton of this approach, that can be overcome by targeted methods. Formation of methy-luridine derivates at the wobble position involves the enzymatic activity of MnmE and GidA (MnmG) and in some tRNAs the position is further modified by MnmC. The 5-methylaminomethyl-2-thiouridine (mnm^5^s^2^U) modification, found in *E. coli* tRNAs tRNA^Lys(UUU)^ and tRNA^GltT/Glu1(UUC)^ is conserved in *P. aeruginosa*, and requires activity of all three enzymes. Synthesis of mnm^5^s^2^U responds to growth conditions (Moukadiri et al. 2014) and cells lacking MnmC are not able to compete with a wild type strain in growth assays (Moukadiri et al. 2014). Two particular modifications, m^1^G and m^2^A occur at position 37 and result in strong termination signatures. m^1^G is deposited by TrmD in *E. coli*, an enzyme that is essential in most bacterial species as it ensures accurate translation and prevents premature termination (Jaroensuk et al. 2019; Björk et al. 2001; Hagervall et al. 1993; Li and Björk 1995; Björk et al. 1989). Reduced levels of m^1^G impair membrane structure and sensitize cells to antibiotics (Masuda et al. 2019), suggesting a possible role for m^1^G in antibiotic resistance. m^2^A is deposited by RlmN (Benítez-Páez et al. 2012; Fitzsimmons and Fujimori 2016). The majority of m^1^G and m^2^A sites had conserved signatures between *P. aeruginosa* and *E. coli*. However, in the anticodon loop of tRNA^His1(GUG)^ and tRNA^Arg1(CCG)^ while we observed a mutational signature at position 37 in both species, the nucleotide identity at this position is different, indicating that different modifications are present (Figure 3 and Supplemental Table 7). In both cases, a strong termination is observed.

### Variable loop modifications in P. aeruginosa

In the variable loop there are two known modifications in *E. coli*, m^7^G deposited at position 46 by YggH and acp^3^U deposited at position 47 by TapT (Takakura et al. 2019; Meyer et al. 2020). While there is no phenotype associated with loss of YggH in *E. coli*, deletion of TrmB, the enzyme responsible for m^7^G in *P. aeruginosa*, induces increased sensitivity to H_2_O_2_ (Thongdee et al. 2019). Levels of acp^3^U are influenced by the presence of m^7^G and culture conditions (Meyer et al. 2020). While one study found loss of TapT to impact thermostability of tRNAs, cell motility and genome stability (Takakura et al. 2019) a different study found no impact of loss of TapT/YfiP on growth under different culture conditions (Meyer et al. 2020). The lack of mutational signature in tRNAs annotated to be 5’-m^7^G U -3’ and the observation that RT stops are absent at position 47 in PA14_17650 mutant cells suggests that acp^3^U is the main contributor for the mutational signature observed at positions 46 and 47 (Meyer et al. 2020). We see conservation of the mutational signature in four tRNAs, with two other sites not showing evidence of being modified in *P. aeruginosa*, despite conservation of the GU sequence at positions 46 and 47. Signatures unique to *P. aeruginosa* are observed in tRNA^Arg1(CCG)^, tRNA^fMet1(CAU)^, tRNA^Asn1(GUU)^ and tRNA^Arg1(UCU)^ (Figure 4, Supplemental Table 7). For tRNA^Arg1(UCU)^ the U required for acp^3^U is absent in *E. coli* tRNA^ArgU(UCU)^ but present in *P. aeruginosa* (Supplemental Table 7). Additionally, we observe a mutational signature in tRNA^Gln1(UUG)^ in a sequence context that is different from *E. coli*. In *V. cholerae*, position 46 of the homologous tRNA is acacp^3^U modified by the enzyme AcpA (Kimura et al. 2020). The same enzyme also modifies U20B in tRNA^Glu1(UUC)^, a position where we also observe a mutational signature in *P. aeruginosa* that is absent in the homologous *E. coli* tRNA^GltT(UUC)^ (Figure 1). While the similarities between *P. aeruginosa* and *V. cholerae* suggest acacp^3^U is also present in *P. aeruginosa*, we were not able to identify a clear homolog for AcpA, although a few enzymes with the same acetyltransferase domain are present the *P. aeruginosa* genome. Using the PA14_17650 mutant strain we validated the presence of acp^3^U on several tRNA in *P. aeruginosa* (Supplemental Table 12, Figure 5B), including tRNA^Gln1(UUG)^. However, we did not observe loss of signature in the PA14_17650 mutant for tRNA^Glu1(UUC)^, tRNA^fMet1(CAU)^, and tRNA^fMet2(CAU)^. There are at least two possible explanations for this observation. Either these positions are modified by a different enzyme with a DTW domain (Takakura et al. 2019), or a unrelated modification is present at this position..

Here we leveraged the wealth of information on *E. coli* tRNAs to explore the landscape of tRNA modifications on *P. aeruginosa*. With this approach we were able to infer which tRNA modifications are conserved in the two species. Our data supports a high degree of conservation between the two organisms and suggests the presence of tRNA modifications in *P. aeruginosa* that are not present in *E. coli*. Determining the specific modifications on each tRNA will provide insights into which pathways are impacted by different tRNA modifying enzymes. Altogether, the approach described here can be used to study tRNA modifications in other unexplored species, or across different culture conditions. Identifying RNA modifications and enzymatic pathways can help reveal targets to potentially reduce pathogenicity, virulence, and antibiotic resistance.

## Materials and Methods

### Strains used in this work

*Escherichia coli* wild type (BW25113) and *E. coli* ΔthiI (JW0413-1, KEIO collection); *Pseudomonas aeruginosa* (PA14). Bacterial strains were grown in liquid media at 37°C and shaken at 235 rpm. For all experiments, bacterial strains were inoculated in LB overnight. Cultures were diluted 1:100 or in the indicated media, grown in triplicate and collected at (OD)_600_=0.6 (log phase). Chemically defined media (CDM) was prepared as described in (Khare and Tavazoie 2015). For the *E. coli* cultures grown in chemically defined media, cultures were diluted 1:250. *Pseudomonas aeruginosa* was grown in CDM. Media used to culture cells in this study: LB (Luria Bertoni, KD Medical; BLF-7030); chemically defined media [1X M63 salts + 0.4 mM ferric citrate + Biotin (0.1 mg/mL) + Nicotinic acid (2 mg/mL) + 1x EZ Supplement + 1x ACGU + 1mM MgSO_4_ + Glucose (0.3%)].

### RNA isolation and RNA library preparation

RNA was isolated with Trizol reagent (ThermoFisher #15596018) (1.2 mL per 25 mL culture after pelleting cells). Samples were extracted with 240 *μ*l of chloroform (Fisher C298-500). Precipitation of the aqueous layer was accompanied with 1 *μ*l of 0.1 M DTT (18091050) and 15 *μ*g of GlycoBlue reagent (ThermoFisher #AM9516). Purified RNA was subject to a second round of isolation using Qiagen RNA columns (74106) with 50 *μ*l RNA + 3.5 volumes of RLT buffer + 10.5 volumes of 100% ethanol. RNA was washed twice following the manufacturers protocol, and eluted 2x in 50 ml of RNase free water. 25 *μ*g of RNA was incubated with 7.5 *μ*g of biotin-conjugated MTSEA (1 mg/ml) (90066-1) in MTSEA buffer [20 mM HEPES pH 7.3 (A611-J848-10), 10 mM EDTA (AM9261)] at 37°C for 2.5 hrs protected from light (600 rpm shake). After labeling with MTSEA, RNA was cleaned again on Qiagen RNA columns. Labeling RNA with MTSEA results in a reversible covalent link by nucleophilic disulfide exchange between sulfur on s^4^U and methanethiosulfonate. The now biotinylated s^4^U-containing RNAs were then captured using C1 streptavidin beads (65001). 25 *μ*l of C1 beads per sample were prepared by washing twice with 500 ml of water, followed by 2 washes in high salt buffer [100 mM Tris pH 7.4 (RGF-3340), 10 mM EDTA (AM9261), 1 M NaCl (AM9759), 0.05% Tween-20 (85113)]. 11 *μ*l of high salt buffer was added to each RNA sample and combined with beads resuspended in 11 *μ*l of high salt buffer. RNA and beads were tumbled in the dark for 30 minutes at room temperature. After capture, beads containing s^4^U-RNA were washed three times in high salt buffer, followed by three washes in NT2 buffer [50 mM Tris pH 7.4 (RGF-3340), 1 mM MgCl_2_ (AM9530G), 150 mM NaCl (AM9759), 0.0005% NP-40 (11332473001)]. Following washes, s^4^U-containing RNAs were eluted by tumbling for 15 min in the dark at room temperature in 100 ml elution buffer [100 mM DTT (18091050), 20 mM HEPES pH 7.3 (A611-J848-10), 1mM EDTA (AM9261), 100 mM NaCl (AM9759), 0.05% Tween-20 (85113)]. Elution was repeated one more time, and the volumes from both extractions combined. RNA was cleaned using the Zymo RNA Clean and Concentrator kit (R1016) following the manufacturers protocol and eluted twice in 7.5 *μ*l of H_2_O.

Sequencing libraries were prepared from total RNA and captured s^4^U-containing RNA. 3^*′*^ end adaptors (Supplemental Table 14) were ligated to 100 ng of RNA in an overnight reaction at 16° C [250 nM adaptor, 1x T4 RNA ligase buffer, 1 *μ*l of T4 RNA Ligase 1 (ssRNA Ligase), High Concentration (M0437M), 50 mM DTT (Sigma 43816-10mL), 6 *μ*l of 50% PEG 8000 (NEB B1004A), 40 units of Ribolock RNase inhibitor EO0382)]. All samples were then gel purified on a denaturing 10% polyacrylamide gel (National Diagnostics, EC-829). Gels were cut between 60 to 200 nt and purified via crush and soak overnight tumble at 4° C [crush gel through 18 gauge needle punched bottom of 500 *μ*l tubes into 1.7 mL Eppendorf tubes, high speed spin for 10 min at RT, and soak crushed gel in 10 mM Tris pH 7.5 (15567-027), 500 mM NaCl (AM9759), 1 mM EDTA (AM9261), 0.1% SDS (351-032-101)]. Gel buffer mix was then transferred to SpinX columns (CLS8162-96EA) to remove remaining gel. Three volumes of 100% ethanol were added to the RNA-containing buffer for overnight precipitation. The purified RNA was then reverse transcribed using TGIRT and uniquely barcoded RT oligonucleotides per sample. Oligonucleotide design and library preparation was adapted from a method previously described (Zarnegar et al. 2016) (Supplemental Table 14). RT included the RNA plus 1 *μ*l of TGIRT enzyme (TGIRT50 from InGex), 1x buffer (20 mM Tris HCl pH 7.5 (15567-027), 5 mM MgCl_2_ (AM9530G), 450 mM NaCl (AM9759)), 1 *μ*l 100 mM DTT (18091050), and 1 *μ*l of 3 mM RT oligonucleotide, in a 19 *μ*l reaction. This mix was incubated for 30 min at room temperature, followed by the addition of 1 *μ*l of 25 mM dNTPs (N0446S), and incubation for 1 hour at 60° C. cDNAs were then released from the RT TGIRT complex by treating with 1 *μ*l of 5M NaOH, incubating at 95° C for 3 min, and followed by neutralization of cooled sample with 1 *μ*l of 5M HCl. cDNAs were then pooled in groups, such as inputs or captured samples, and cleaned using the Zymo DNA Clean and Concentrator kit (D4014). Individually barcoded and cleaned pools of samples were then gel purified to remove excess oligonucleotide, following the same protocol as above, except that eluted cDNA was cleaned through Zymo DNA Clean and Concentrator kit (D4014). Eluted pools were then subjected to circularization using a CircLigase-II kit (CL9021K) (cDNA, 0.75 *μ*l MnCl_2_, 0.75 *μ*l CircLigase-II, 1.5 *μ*l of 10x buffer in a 15 *μ*l reaction). After circularization, cDNA was cleaned mixing with 30 ml of Ampure XP beads (A63881) and 75 *μ*l of isopropanol, incubated 10 min RT, applied to magnet, washed two times 30 seconds each with freshly made 80% EtOH. Beads with cDNA were air dried for 5 min and cDNA was then eluted from beads in 14 *μ*l of H_2_O. Directly following clean up, cDNA was amplified with a first set of oligos in a PCR reaction on a qPCR machine to detect amplification in real time to capture linear amplification (0.75 *μ*l of 20 mM P3^tall^ and P6^tall^ oligo mix, 15 *μ*l of 2x NEB Next (M0541L), 14 *μ*l cDNA, and 0.25 *μ*l 25x diluted Sybr green (S7563) in a 30 *μ*l reaction). PCR parameters were 98° C for 40 seconds, cycling of 98° C for 20 seconds, 65° C for 30 seconds, 72° C for 45 seconds, and a final extension at 72° C for 5 minutes. Number of cycles was determined by real time amplification. Samples were then cleaned again using Ampure XP beads (A63881) (54 *μ*l of beads, 4 *μ*l of isopropanol, and eluted in *μ*l ml of H_2_O). Eluted cDNA was amplified for three more cycles using a second set of oligonucleotides for Illumina sequencing in a NEB Next master mix reaction. Following the second PCR, samples were cleaned one last time on the Ampure XP beads (36 *μ*l of beads, 2.7 *μ*l of isopropanol, and eluted in 12 *μ*l of H_2_O). Final libraries were analyzed by DNA high sensitivity tape station.

### Targeted RT sequencing

Adapted from a previously described method we performed targeted RT sequencing (Link et al. 2022). Briefly, RNA was isolated from Trizol Reagent (ThermoFisher #15596018) with chloroform according to the manufacturer’s protocol, except that RNA was isolated from the aqueous layer using Qiagen columns, washed 2x in 500 ml of RPE wash buffer, dried, and eluted with 100 *μ*l of H_2_O. Reverse transcription was performed with 500 ng RNA each using TGIRT (obtained from Lambowitz lab) (Mohr et al. 2013), SuperScript III (ThermoFisher #18080044), SuperScript IV (ThermoFisher #18090050) or Induro (NEB #M0681L) reverse transcriptases. Each RT was performed using an RT oligo that was specifically designed to target the 3^*′*^ end of tRNA^Arg1(ACG)^ (Supplemental Table 14). TGIRT RT was performed just as described in the library preparation, SuperScript III RT was performed following the manufacturer’s protocol (500 ng RNA, 1 *μ*l of 3 mM RT oligo, 4 *μ*l of 5x FS buffer and up to 16 *μ*l total with H_2_O). Samples were annealed at 65° C for 5 min, ice for 1 min, and then supplemented with 1 *μ*l 10 mM dNTPs, 1 *μ*l of 0.1 M DTT, 1 *μ*l RNasin (Promega #N2611), and 1 *μ*l of SuperScript III enzyme, and then incubated at 55° C for 1 hour, followed by 70° C for 15 min. Similarly, SuperScript IV RT was performed following the manufacturer’s protocol. First step (500 ng RNA, 1 *μ*l of 3 mM oligo, 1 *μ*l of 10 mM dNTP mix and up to 13 *μ*l of H_2_O) incubated at 65° C for 5 min, ice for 1 min, followed by addition of 4 *μ*l of 5x SSIV buffer, 1 *μ*l of 0.1M DTT, 1 *μ*l of RNase inhibitor (ThermoFisher #EO0381), and 1 *μ*l of SSIV enzyme, and incubation at 55° C for 10 min, followed by heat inactivation at 80° C for 10 min. Finally, Induro RT was also performed according to the manufacturer’s protocol first incubating at 65° C for 5 min, and then on ice for 1 min (500 ng RNA, 1 *μ*l of 3 mM oligo, 1 *μ*l of 10 mM dNTP mix up to 10 *μ*l of H_2_O), followed by addition of 4 *μ*l of 5x Induro RT reaction buffer, 0.2 *μ*l of RNase inhibitor, 1 *μ*l of Induro enzyme and 4.8 *μ*l of H_2_O, incubating for 1 hour at 60° C, and heat inactivated at 95° C for 1 min. Following all RT reactions, samples were PCR amplified using HF Phusion enzyme (NEB #M0530L) (2 *μ*l of cDNA, 10 *μ*l of 5x HF buffer, 2.5 *μ*l of 10 mM forward oligo, 2.5 *μ*l of 10 mM reverse RT oligo, 1 *μ*l of 10 mM dNTP, 0.5 *μ*l of Phusion, and 31.5 *μ*l of H_2_O. PCR parameters were 95° C for 1 min, cycling of 95° C for 15 seconds, 61° C for 15 seconds, 72° C for 30 seconds, for 33 cycles, and a final extension at 72° C for 7 minutes. Following the PCR, samples were cleaned with Zymo DNA Clean and Concentrator kit (D4014) following the manufacturer’s protocol and eluted 2x in 11 *μ*l of H_2_O each. Samples were quantified and then submitted for Sanger sequencing using the forward sequencing oligo (oligos listed in Supplemental Table 14).

### tRNA purifications and mass spectrometry

tRNAs were gel purified from total RNA on 10% UREA-polyacrylamide gels. Briefly, 10 to 20 *μ*g of RNA containing 2x RNA loading dye (ThermoFisher #R0641) were heated for 1 min at 95° C before loading onto a pre-run 10% UREA-PAGE gel, in comparison to low molecular weight ssRNA (NEB #N0364S). Gels were cut on a blue light box corresponding to the size of tRNAs (70 to 100 nt) and purified via crush and soak overnight tumble at 4° C [crush gel through 18 gauge needle punched bottom of 500 *μ*l tubes into 1.7 mL Eppendorf tubes, high speed spin for 10 min at RT, and soak crushed gel in 10 mM Tris pH 7.5 (15567-027), 500 mM NaCl (AM9759), 1 mM EDTA (AM9261), 0.1% SDS (351-032-101)]. Gel buffer mix was then transferred to SpinX columns (Sigma #CLS8162-96EA) to remove remaining gel, and 1 *μ*lof GlycoBlue co-precipitant (ThermoFisher #AM9516). Three volumes of 100% ethanol were added to the RNA-containing buffer for overnight precipitation. Ethanol precipitations were then centrifuged at high speed for 45 min at 4° C, washed 1x with 75% ethanol, air dried for 5 min, and resuspended in 50 *μ*l of H_2_O. Purified tRNAs were then re-cleaned using Zymo RNA Clean and Concentrator kit (R1016) according to the manufacturer’s protocol, except we used three volumes of 100% ethanol in the first step. Samples were eluted 2x 11 *μ*l of H_2_O and quantified by DeNovix spectrophotometer (DS-11+) before preparing for mass spectrometry.

Total tRNA or decapped RNA was subsequently digested to nucleosides in a 30 *μ*l reaction containing 24 *μ*l RNA, 1 *μ*l of 2U/*μ*l Nuclease P1 (Sigma Aldrich, #N8630), 3 *μ*l 1M ammonium acetate, pH 5.2, and 100 fmol of ^13^C-adenosine (Cambridge Isotope #CLM-3678-0.05) as an internal standard. Reactions were digested for 3 hr at 45° C. After 3 hours, 3*μ*l of 1M ammonium bicarbonate and 1 *μ*l of 0.002 U/*μ*l of phosphodiesterase from *Crotalus ademanteus* (Sigma Aldrich #P3243) were added to the reaction and incubation continued for 2 hrs at 37° C. After 2 hrs, 1 *μ*l of 1U/*μ*l bacterial alkaline phosphatase (ThermoFisher #18011015) was added to the reaction and the reaction continued for 1 hr at 37° C. Digested nucleosides were passed through a 3 kDa MWCO spin filter (Amicon # UFC500396) and the filter was rinsed with 3 × 200 *μ*l nuclease free water before being dried under vacuum. Dried samples were dissolved in 45 *μ*l Buffer A (0.1% formic acid). Nucleosides were separated on a Luna Omega C18 reverse-phase column (1.6 mm PS, 100 Å, 30 mm, ID 2.1 mm, Phenomenex) with a flow rate of 0.200 *μ*l/min under the following conditions. Buffer A = 0.1% formic acid in high performance liquid chromatography (HPLC)-grade water; Buffer B = 0.1% formic acid in acetonitrile. The gradient for the liquid chromatography is as follows: Gradient 0.2% Buffer B from 0-2.6 min; Gradient 2% Buffer B from 2.6-5.0 min; Gradient 8.5% B from 5.01-6.75 min; Gradient 40% B from 6.76-7.5 min. 50% Buffer B from 7.51-10 min; 0% Buffer B from 10.01-14.0 min. Following chromatography, nucleosides were resolved on a SciEX 500 QTOF in positive electrospray ionization mode using a Multiple-reaction-monitoring (MRM) MS method over the range of 100-500 daltons (Spray voltage = 5500 V; Curtain gas = 35; CAD gas = 7; Temp = 500° C; Declustering potential = 25 V; Collision energy = 10 V; Accumulation time = 0.25 sec). The nucleosides were quantified using the nucleoside to base ion mass transitions. Chemical standards for the modified acp^3^U were obtained from Smolecule (S562863).

### Modifying enzyme homology search between P. aeruginosa and E. coli

Enzymes responsible for tRNA modifications in *E. coli* were identified from a search of the literature (Kimura et al. 2020; de Crécy-Lagard and Jaroch 2021). Accession numbers, genomic loci, and corresponding peptide sequences were retrieved from NCBI’s gene lookup tool, specifically referencing *E. coli* K-12 MG1655 (NCBI:txid511145). Peptide sequences were used to carry out a BLASTP search using the *Pseudomonas* Genome Database’s tool looking for proteins found in the *P. aeruginosa* strain PA14 (UCBPP-PA14 2006) (Winsor et al. 2016). If a match was found, the top hit was listed along with the total number of hits. Homologous proteins are listed in Supplemental Table 12.

### RNA sequencing analysis

RNA sequencing was performed using NextSeq 550 or 2000 Illumina sequencers. Sequence reads were separated by barcode and duplicated reads removed. Adapters sequences were trimmed. Reads were mapped with Bowtie2 to a custom transcriptome using a local alignment tool. Read counts were compiled for each sample and analyzed using DESeq2 to identify enrichment of reads from s^4^U-captured RNAs as compared to inputs (Supplemental Tables 3 and 8). tRNAs were considered enriched if l2fc > 0.6, padj < 0.05, and > 100 read coverage in the input sample. To generate custom transcriptomes for both *E. coli* and *P. aeruginosa*, fasta files for annotated cDNA and ncRNA, were downloaded from Ensembl for genome assembly ASM584v2 (GCA_000005845) and ASM1462v1 (GCA_000014625.1), respectively. For tRNA genes, duplicated sequences were removed, so that only unique sequences are present in the final transcriptome. CCA was added to the 3^*′*^ end on tRNAs in the *P. aeruginosa* transcriptome. Codon identity for each *P. aeruginosa* tRNA gene was obtained from the tRNA database (Jühling et al. 2009). Annotated modification information was obtained from (Boccaletto et al. 2022; Kimura et al. 2020). For all misincorporation and early termination analyses, only positions with 100 reads or higher were considered. *E. coli* and *P. aeruginosa* tRNA were matched based on anticodon and by sequence similarity, determined with the Emboss Needle tool (https://www.ebi.ac.uk/Tools/psa/emboss_needle/). To determine conservation between tRNA modifying enzymes we used a protein Blast tool from www.pseudomonas.com, using *E. coli* protein sequences obtained from https://www.ncbi.nlm.nih.gov/protein/.

### Dot blot and acrylamide gel analysis

5 mg of RNA was incubated with 2.5 mg biotin-conjugated MTSEA (1 mg/ml) (Biotium 90066-1) in MTSEA buffer [20 mM HEPES pH 7.3 (Quality Biological A611-J848-10), 10 mM EDTA (Invitrogen AM9261] at 37°C for 2.5 hrs (600 rpm shake). Samples were extracted with 100 *μ*l of chloroform (Fisher C298-500) and the aqueous phase was processed on a Zymo column (Zymo R1018). 300 ng of each sample was loaded on Hybond-N^+^ membrane (Amersham RPN119B). Membrane was crosslinked with UV (254 nm) for 120000 mJoules, twice. Membrane was blocked in Odyssey blocking buffer (LiCor 927-40003) for 10 min. IRDye® 800CW Streptavidin (LiCor P/N 926-32230) was added to blocking buffer at 1:1000 dilution and incubated for 45 min. Membrane was washed with Phosphate Buffer Saline (PBS) (pH 7.2) (KD Medical) Tween 20 (0.1%) (Sigma P9416) two times and once with PBS (KD Medical) and imaged on ODYSSEY CLx (LiCor). A solution of 0.04% Methylene blue (Sigma M4159) in 0.5M NaOAc (Invitrogen AM9740) was used to stain nucleic acids for loading control. Membrane was washed with nuclease-free water and imaged on a ChemiDoc MP Imaging system (BioRad). Denaturing polyacrylamide gels were made using the SequaGel UreaGel 29:1 Denaturing Gel System (EC-829). Non-denaturing gels were made with acrylamide (30% Acrylamide/Bis Solution, 29:1 (1610156)), 1xTBE (BP13334), 0.1% ammonium persulfate, and 0.1% TEMED (1610800). Gels were run with size standards for RNA or DNA [low range ssRNA ladder (N0364S); low molecular weight DNA ladder (N3233L)]. Gels were visualized by staining with Sybr Gold nucleic acid gel stain (S11494).

### Data Availability and Supplemental Tables

The raw sequencing data reported in this paper have been deposited in the Gene Expression Omnibus (GEO) at GSE236676. Supplemental tables, as well as code and scripts used to analyze the data and generate the figures are available at https://github.com/BatistaLab/Prokaryotic_tRNA.

## Abbreviations

(LB): Luria-Bertani
(CDM): Chemically defined media
(UV): Ultraviolet
(*E. coli*): *Escherichia coli*
(P. aeruginosa): Pseudomonas aeruginosa
(s^4^U): 4-thiouridine
(acp^3^U): 3-(3-amino-3-carboxypropyl)uridine
(acacp^3^U): acetylated aminocarboxypropyluridine
(cmo^5^U): Uridine 5-oxyacetic acid
(m^5^U): 5-methyluridine
(mnm^5^s^2^U): 5-methylaminomethyl-2-thiouridine
(Q): Queuosine
(Y): Pseudouridine
(mcmo^5^U): Uridine 5-oxyacetic acid methyl ester
(ac^4^C): N4-acetylcytidine
(m^1^G): 1-methylguanosine
(Um): 2^*′*^-O-methyluridine
(ncm^5^s^2^U): 5-carbamoylmethyl-2-thiouridine
(m^6^t^6^A): N6-methyl-N6-threonylcarbamoyladenosine
(D): Dihydrouridine
(Cm): 2^*′*^-O-methylcytidine
(m^7^G): 7-methylguanosine
(ct^6^A): cyclic N6-threonylcarbamoyladenosine
(cmnm^5^s^2^U): 5-carboxymethylaminomethyl-2-thiouridine
(gluQ): Glutamyl-queuosine
(m^6^A): N6-methyladenosine
(s^2^C): 2-thiocytidine
(Gm): 2^*′*^-O-methylguanosine
(m^2^A): 2-methyladenosine
(ms^2^i^6^A): 2-methylthio-N6-isopentenyladenosine
(k^2^C): 2-lysidine
(mnm^5^U): 5-methylaminomethyluridine
(cmnm^5^Um): 5-carboxymethylaminomethyl-2^*′*^-O-methyluridine

## ACKNOWLEDGEMENTS

We thank Susan Gottesman, Richard Maraia and members of Batista lab for critical reading of the manuscript, as well as Anupama Khare for providing bacteria, reagents and protocols. We also thank Alan Lambowitz for providing us with TGIRT enzyme. The authors thank the Center for Cancer Research (CCR) Genomics Core in Bethesda, Maryland for help with high-throughput sequencing. This work utilized the computational resources of the NIH HPC Biowulf cluster (http://hpc.nih.gov). We appreciate the editorial assistance of George Leiman. Research was supported by the Intramural Research Program at the National Cancer Institute (NCI) of the National Institutes of Health.

## Bibliography

Benítez-Páez A, Villarroya M, Armengod M-E. 2012. The Escherichia coli RlmN methyltransferase is a dual-specificity enzyme that modifies both rRNA and tRNA and controls translational accuracy. RNA 18: 1783–1795.

Björk GR, Hagervall TG. 2014. Transfer RNA Modification: Presence, Synthesis, and Function. EcoSal Plus 6.

Björk GR, Jacobsson K, Nilsson K, Johansson MJ, Byström AS, Persson OP. 2001. A primordial tRNA modification required for the evolution of life? EMBO J 20: 231–239.

Björk GR, Wikström PM, Byström AS. 1989. Prevention of translational frameshifting by the modified nucleoside 1-methylguanosine. Science 244: 986–989.

Boccaletto P, Stefaniak F, Ray A, Cappannini A, Mukherjee S, Purta E, Kurkowska M, Shirvanizadeh N, Destefanis E, Groza P, et al. 2022. MODOMICS: a database of RNA modification pathways. 2021 update. Nucleic Acids Res 50: D231–D235.

Bommisetti P, Bandarian V. 2022. Site-Specific Profiling of 4-Thiouridine Across Transfer RNA Genes in Escherichia coli. ACS Omega 7: 4011–4025.

Cho KH, Caparon MG. 2008. tRNA modification by GidA/MnmE is necessary for Streptococcus pyogenes virulence: a new strategy to make live attenuated strains. Infect Immun 76: 3176–3186.

Cozen AE, Quartley E, Holmes AD, Hrabeta-Robinson E, Phizicky EM, Lowe TM. 2015. ARM-seq: AlkB-facilitated RNA methylation sequencing reveals a complex landscape of modified tRNA fragments. Nat Methods 12: 879–884.

de Crécy-Lagard V, Jaroch M. 2021. Functions of Bacterial tRNA Modifications: From Ubiquity to Diversity. Trends Microbiol 29: 41–53.

de Crécy-Lagard V, Ross RL, Jaroch M, Marchand V, Eisenhart C, Brégeon D, Motorin Y, Limbach PA. 2020. Survey and Validation of tRNA Modifications and Their Corresponding Genes in Bacillus subtilis sp Subtilis Strain 168. Biomolecules 10: 977.

Duechler M, Leszczyńska G, Sochacka E, Nawrot B. 2016. Nucleoside modifications in the regulation of gene expression: focus on tRNA. Cell Mol Life Sci 73: 3075–3095.

Duffy EE, Rutenberg-Schoenberg M, Stark CD, Kitchen RR, Gerstein MB, Simon MD. 2015. Tracking Distinct RNA Populations Using Efficient and Reversible Covalent Chemistry. Mol Cell 59: 858–866.

Dunin-Horkawicz S, Czerwoniec A, Gajda MJ, Feder M, Grosjean H, Bujnicki JM. 2006. MODOMICS: a database of RNA modification pathways. Nucleic Acids Res 34: D145–149.

Durand JM, Dagberg B, Uhlin BE, Björk GR. 2000. Transfer RNA modification, temperature and DNA superhelicity have a common target in the regulatory network of the virulence of Shigella flexneri: the expression of the virF gene. Mol Microbiol 35: 924–935.

Emilsson V, Näslund AK, Kurland CG. 1992. Thiolation of transfer RNA in Escherichia coli varies with growth rate. Nucleic Acids Res 20: 4499–4505.

Esberg B, Leung HC, Tsui HC, Björk GR, Winkler ME. 1999. Identification of the miaB gene, involved in methylthiolation of isopentenylated A37 derivatives in the tRNA of Salmonella typhimurium and Escherichia coli. J Bacteriol 181: 7256–7265.

Favre A, Michelson AM, Yaniv M. 1971. Photochemistry of 4-thiouridine in Escherichia coli transfer RNA1Val. J Mol Biol 58: 367–379.

Fitzsimmons CM, Fujimori DG. 2016. Determinants of tRNA Recognition by the Radical SAM Enzyme RlmN. PLoS One 11: e0167298.

Flynn RA, Zhang QC, Spitale RC, Lee B, Mumbach MR, Chang HY. 2016. Transcriptome-wide interrogation of RNA secondary structure in living cells with icSHAPE. Nat Protoc 11: 273–290.

Frye M, Harada BT, Behm M, He C. 2018. RNA modifications modulate gene expression during development. Science 361: 1346–1349.

Gogakos T, Brown M, Garzia A, Meyer C, Hafner M, Tuschl T. 2017. Characterizing Expression and Processing of Precursor and Mature Human tRNAs by Hydro-tRNAseq and PAR-CLIP. Cell Rep 20: 1463–1475.

Grobe S, Doberenz S, Ferreira K, Krueger J, Brönstrup M, Kaever V, Häussler S. 2019. Identification and Quantification of (t)RNA Modifications in Pseudomonas aeruginosa by Liquid Chromatography-Tandem Mass Spectrometry. Chembiochem 20: 1430–1437.

Gupta R, Gobble TR, Schuster M. 2009. GidA posttranscriptionally regulates rhl quorum sensing in Pseudomonas aeruginosa. J Bacteriol 191: 5785–5792.

Hagervall TG, Tuohy TM, Atkins JF, Björk GR. 1993. Deficiency of 1-methylguanosine in tRNA from Salmonella typhimurium induces frameshifting by quadruplet translocation. J Mol Biol 232: 756–765.

Jaroensuk J, Atichartpongkul S, Chionh YH, Wong YH, Liew CW, McBee ME, Thongdee N, Prestwich EG, DeMott MS, Mongkolsuk S, et al. 2016. Methylation at position 32 of tRNA catalyzed by TrmJ alters oxidative stress response in Pseudomonas aeruginosa. Nucleic Acids Res 44: 10834–10848.

Jaroensuk J, Wong YH, Zhong W, Liew CW, Maenpuen S, Sahili AE, Atichartpongkul S, Chionh YH, Nah Q, Thongdee N, et al. 2019. Crystal structure and catalytic mechanism of the essential m1G37 tRNA methyltransferase TrmD from Pseudomonas aeruginosa. RNA 25: 1481–1496.

Jühling F, Mörl M, Hartmann RK, Sprinzl M, Stadler PF, Pütz J. 2009. tRNAdb 2009: compilation of tRNA sequences and tRNA genes. Nucleic Acids Res 37: D159–162.

Khare A, Tavazoie S. 2015. Multifactorial Competition and Resistance in a Two-Species Bacterial System. PLoS Genet 11: e1005715.

Kimura S, Dedon PC, Waldor MK. 2020. Comparative tRNA sequencing and RNA mass spectrometry for surveying tRNA modifications. Nat Chem Biol 16: 964–972.

Kimura S, Miyauchi K, Ikeuchi Y, Thiaville PC, Crécy-Lagard V de, Suzuki T. 2014. Discovery of the β-barrel-type RNA methyltransferase responsible for N6-methylation of N6-threonylcarbamoyladenosine in tRNAs. Nucleic Acids Res 42: 9350–9365.

Kimura S, Waldor MK. 2019. The RNA degradosome promotes tRNA quality control through clearance of hypomodified tRNA. Proc Natl Acad Sci U S A 116: 1394–1403.

Kinscherf TG, Willis DK. 2002. Global regulation by gidA in Pseudomonas syringae. J Bacteriol 184: 2281–2286.

Koh CS, Sarin LP. 2018. Transfer RNA modification and infection - Implications for pathogenicity and host responses. Biochim Biophys Acta Gene Regul Mech 1861: 419–432.

Kramer GF, Baker JC, Ames BN. 1988. Near-UV stress in Salmonella typhimurium: 4-thiouridine in tRNA, ppGpp, and ApppGpp as components of an adaptive response. J Bacteriol 170: 2344–2351.

Langmead B, Salzberg SL. 2012. Fast gapped-read alignment with Bowtie 2. Nat Methods 9: 357–359.

Li JN, Björk GR. 1995. 1-Methylguanosine deficiency of tRNA influences cognate codon interaction and metabolism in Salmonella typhimurium. J Bacteriol 177: 6593–6600.

Liberati NT, Urbach JM, Miyata S, Lee DG, Drenkard E, Wu G, Villanueva J, Wei T, Ausubel FM. 2006. An ordered, nonredundant library of Pseudomonas aeruginosa strain PA14 transposon insertion mutants. Proc Natl Acad Sci U S A 103: 2833–2838.

Lin Q, Huang J, Liu Z, Chen Q, Wang X, Yu G, Cheng P, Zhang L-H, Xu Z. 2022. tRNA modification enzyme MiaB connects environmental cues to activation of Pseudomonas aeruginosa type III secretion system. PLoS Pathog 18: e1011027.

Link CN, Thalalla Gamage S, Gallimore D, Kopajtich R, Evans C, Nance S, Fox SD, Andresson T, Chari R, Ivanic J, et al. 2022. Protonation-Dependent Sequencing of 5-Formylcytidine in RNA. Biochemistry 61: 535–544.

Lyczak JB, Cannon CL, Pier GB. 2000. Establishment of Pseudomonas aeruginosa infection: lessons from a versatile opportunist. Microbes Infect 2: 1051–1060.

Masuda I, Matsubara R, Christian T, Rojas ER, Yadavalli SS, Zhang L, Goulian M, Foster LJ, Huang KC, Hou Y-M. 2019. tRNA Methylation Is a Global Determinant of Bacterial Multi-drug Resistance. Cell Syst 8: 302-314.e8.

Meyer B, Immer C, Kaiser S, Sharma S, Yang J, Watzinger P, Weiß L, Kotter A, Helm M, Seitz H-M, et al. 2020. Identification of the 3-amino-3-carboxypropyl (acp) transferase enzyme responsible for acp3U formation at position 47 in Escherichia coli tRNAs. Nucleic Acids Res 48: 1435–1450.

Mohr S, Ghanem E, Smith W, Sheeter D, Qin Y, King O, Polioudakis D, Iyer VR, Hunicke-Smith S, Swamy S, et al. 2013. Thermostable group II intron reverse transcriptase fusion proteins and their use in cDNA synthesis and next-generation RNA sequencing. RNA 19: 958–970.

Moukadiri I, Garzón M-J, Björk GR, Armengod M-E. 2014. The output of the tRNA modification pathways controlled by the Escherichia coli MnmEG and MnmC enzymes depends on the growth conditions and the tRNA species. Nucleic Acids Res 42: 2602–2623.

Moukadiri I, Prado S, Piera J, Velázquez-Campoy A, Björk GR, Armengod M-E. 2009. Evolutionarily conserved proteins MnmE and GidA catalyze the formation of two methyluridine derivatives at tRNA wobble positions. Nucleic Acids Res 37: 7177–7193.

Mueller EG, Buck CJ, Palenchar PM, Barnhart LE, Paulson JL. 1998. Identification of a gene involved in the generation of 4-thiouridine in tRNA. Nucleic Acids Res 26: 2606–2610.

Pan T. 2018. Modifications and functional genomics of human transfer RNA. Cell Res 28: 395–404.

Persson BC, Björk GR. 1993. Isolation of the gene (miaE) encoding the hydroxylase involved in the synthesis of 2-methylthio-cis-ribozeatin in tRNA of Salmonella typhimurium and characterization of mutants. J Bacteriol 175: 7776–7785.

Pritchard JR, Chao MC, Abel S, Davis BM, Baranowski C, Zhang YJ, Rubin EJ, Waldor MK. 2014. ARTIST: high-resolution genome-wide assessment of fitness using transposon-insertion sequencing. PLoS Genet 10: e1004782.

Qian Q, Curran JF, Björk GR. 1998. The methyl group of the N6-methyl-N6-threonylcarbamoyladenosine in tRNA of Escherichia coli modestly improves the efficiency of the tRNA. J Bacteriol 180: 1808–1813.

Qin Y, Yao J, Wu DC, Nottingham RM, Mohr S, Hunicke-Smith S, Lambowitz AM. 2016. High-throughput sequencing of human plasma RNA by using thermostable group II intron reverse transcriptases. RNA 22: 111–128.

Ramabhadran TV, Jagger J. 1976. Mechanism of growth delay induced in Escherichia coli by near ultraviolet radiation. Proc Natl Acad Sci U S A 73: 59–63.

Richter U, Evans ME, Clark WC, Marttinen P, Shoubridge EA, Suomalainen A, Wredenberg A, Wedell A, Pan T, Battersby BJ. 2018. RNA modification landscape of the human mitochondrial tRNALys regulates protein synthesis. Nat Commun 9: 3966.

Romsang A, Duang-Nkern J, Khemsom K, Wongsaroj L, Saninjuk K, Fuangthong M, Vattanaviboon P, Mongkolsuk S. 2018. Pseudomonas aeruginosa ttcA encoding tRNA-thiolating protein requires an iron-sulfur cluster to participate in hydrogen peroxide-mediated stress protection and pathogenicity. Sci Rep 8: 11882.

Schwartz MH, Wang H, Pan JN, Clark WC, Cui S, Eckwahl MJ, Pan DW, Parisien M, Owens SM, Cheng BL, et al. 2018. Microbiome characterization by high-throughput transfer RNA sequencing and modification analysis. Nat Commun 9: 5353.

Sha J, Kozlova EV, Fadl AA, Olano JP, Houston CW, Peterson JW, Chopra AK. 2004. Molecular characterization of a glucose-inhibited division gene, gidA, that regulates cytotoxic enterotoxin of Aeromonas hydrophila. Infect Immun 72: 1084–1095.

Stover CK, Pham XQ, Erwin AL, Mizoguchi SD, Warrener P, Hickey MJ, Brinkman FS, Hufnagle WO, Kowalik DJ, Lagrou M, et al. 2000. Complete genome sequence of Pseudomonas aeruginosa PAO1, an opportunistic pathogen. Nature 406: 959–964.

Su C, Jin M, Zhang W. 2022. Conservation and Diversification of tRNA t6A-Modifying Enzymes across the Three Domains of Life. Int J Mol Sci 23: 13600.

Takakura M, Ishiguro K, Akichika S, Miyauchi K, Suzuki T. 2019. Biogenesis and functions of aminocarboxypropyluridine in tRNA. Nat Commun 10: 5542.

Thomas G, Favre A. 1980. 4-Thiouridine triggers both growth delay induced by near-ultraviolet light and photoprotection. Eur J Biochem 113: 67–74.

Thomas G, Thiam K, Favre A. 1981. tRNA thiolated pyrimidines as targets for near-ultraviolet-induced synthesis of guanosine tetraphosphate in Escherichia coli. Eur J Biochem 119: 381–387.

Thongdee N, Jaroensuk J, Atichartpongkul S, Chittrakanwong J, Chooyoung K, Srimahaeak T, Chaiyen P, Vattanaviboon P, Mongkolsuk S, Fuangthong M. 2019. TrmB, a tRNA m7G46 methyltransferase, plays a role in hydrogen peroxide resistance and positively modulates the translation of katA and katB mRNAs in Pseudomonas aeruginosa. Nucleic Acids Res 47: 9271–9281.

Vandivier LE, Anderson ZD, Gregory BD. 2019. HAMR: High-Throughput Annotation of Modified Ribonucleotides. Methods Mol Biol 1870: 51–67.

Wang J, Toffano-Nioche C, Lorieux F, Gautheret D, Lehmann J. 2021. Accurate characterization of Escherichia coli tRNA modifications with a simple method of deepsequencing library preparation. RNA Biol 18: 33–46.

Winsor GL, Griffiths EJ, Lo R, Dhillon BK, Shay JA, Brinkman FSL. 2016. Enhanced annotations and features for comparing thousands of Pseudomonas genomes in the Pseudomonas genome database. Nucleic Acids Res 44: D646–653.

Wolf J, Gerber AP, Keller W. 2002. tadA, an essential tRNA-specific adenosine deaminase from Escherichia coli. EMBO J 21: 3841–3851.

Zarnegar BJ, Flynn RA, Shen Y, Do BT, Chang HY, Khavari PA. 2016. irCLIP platform for efficient characterization of protein-RNA interactions. Nat Methods 13: 489–492.

Zhang W, Foo M, Eren AM, Pan T. 2022. tRNA modification dynamics from individual organisms to metaepitranscriptomics of microbiomes. Mol Cell 82: 891–906.

Zheng G, Qin Y, Clark WC, Dai Q, Yi C, He C, Lambowitz AM, Pan T. 2015. Efficient and quantitative high-throughput tRNA sequencing. Nat Methods 12: 835–837.

