## Supplemental Figures for "The modification landscape of *P. aeruginosa* tRNAs"

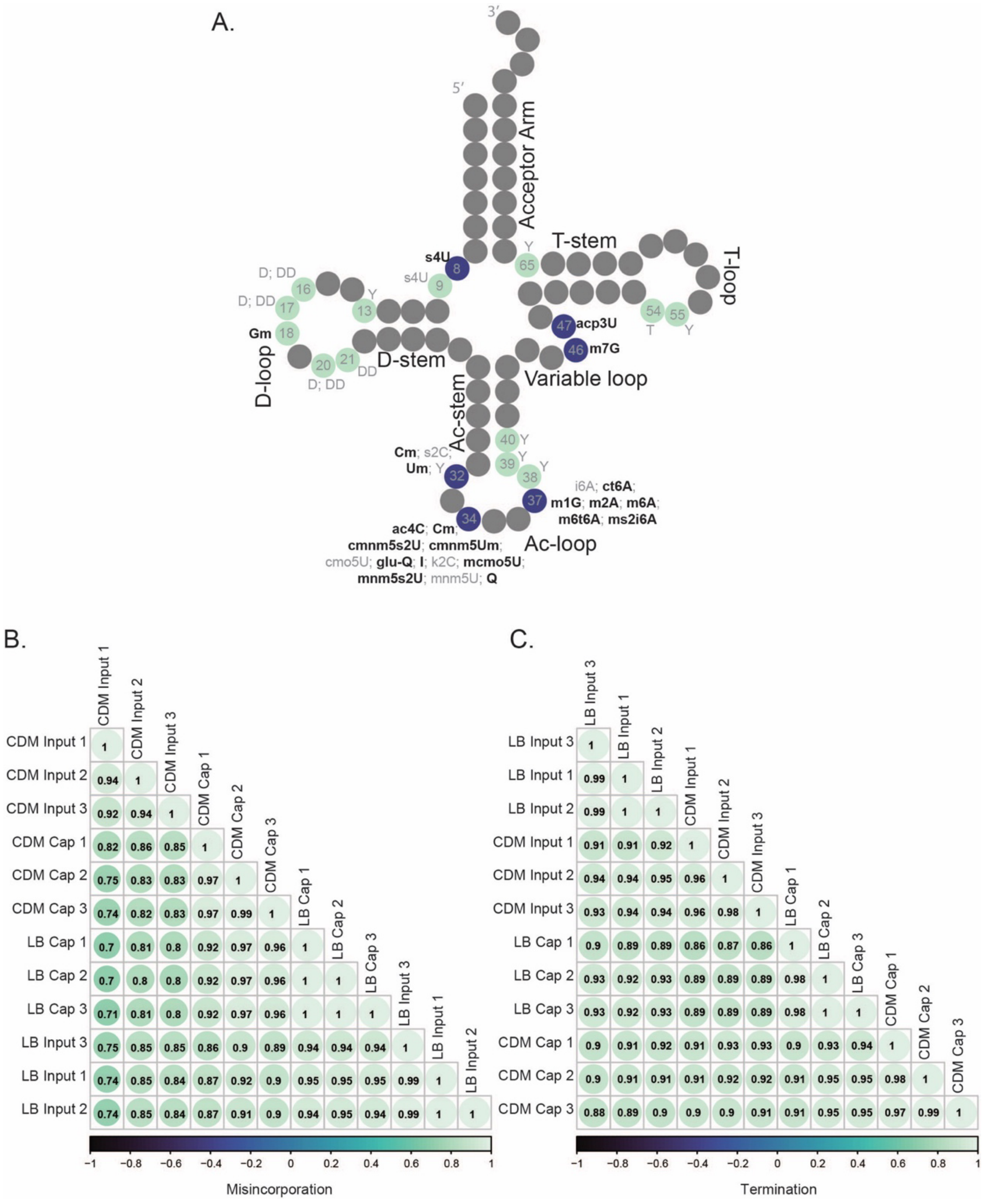


**Supplemental Figure 1.**

**(A)** Schematic representation of a mature tRNA. Positions known to be modified are indicated in green (positions where a cDNA mutational signature is not observed) or blue (positions where a mutational signature is observed). Modifications with a detectable mutational signature are represented in bold. **(B)** Correlation plots for misincorporation and **(C)** termination between replicates for total RNA (input) and MTSEA-captured (Cap) samples in *E. coli* cultured in chemically defined media (CDM) and LB.

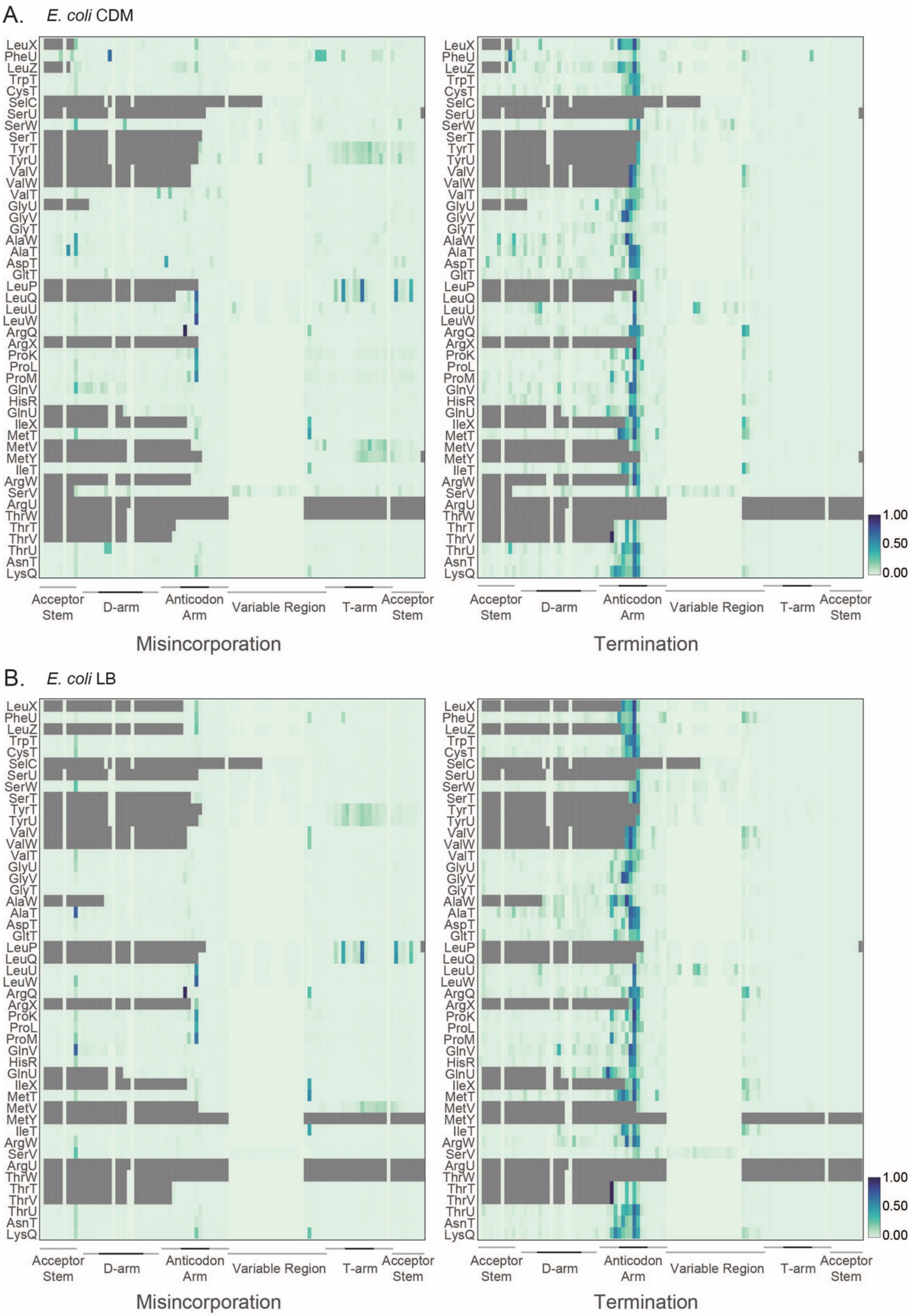
A.
E. coli CDM
LeuX
PheU
LeuZ
TrpT
CysT
SelC
SerU
SerW
SerT
TyrT
TyrU
ValV
ValW
ValT
GlyU
GlyV
GlyT
AlaW
AlaT
AspT
GltT
LeuP
LeuQ
LeuU
LeuW
ArgQ
ArgX
ProK
ProL
ProM
GlnV
HisR
GlnU
IleX
MetT
MetV
MetY
IleT
ArgW
SerV
ArgU
ThrW
ThrT
ThrV
ThrU
AsnT
LysQ
Acceptor Stem
D-arm
Anticodon Arm
Variable Region
T-arm
Acceptor Stem
Misincorporation
Termination
1.00
0.50
0.00
B.
E. coli LB
Misincorporation
Termination
1.00
0.50
0.00

**Supplemental Figure 2.**

**(A)** Heatmap of average frequency (from three independent samples) of misincorporation (left) or termination score (right) at each position for *E. coli* tRNAs extracted from cells cultured in chemically modified media (CDM) and collected at $(OD)_{600}$=0.6. All tRNAs in the custom transcriptome are represented. The color key at the right represents the frequency of misincorporation or termination score. The grey refers to regions of less than 100 reads per position. Different regions of the tRNA are represented at the bottom from the 5′ (left) to the 3′ end (right) of the tRNA. Darker lines indicate the loop in that region. **(B)** Heatmap of average frequency (from three independent samples) of misincorporation (left) or termination score (right) at each position for *E. coli* tRNAs extracted from cells cultured in LB and collected at $(OD)_{600}$=0.6. All tRNAs in the custom transcriptome are represented. The color key at the right represents the frequency of misincorporation or termination score. The grey refers to regions of less than 100 reads per position. Different regions of the tRNA are represented at the bottom from the 5′ (left) to the 3′ end (right) of the tRNA. Darker lines indicate the loop in that region.

A.

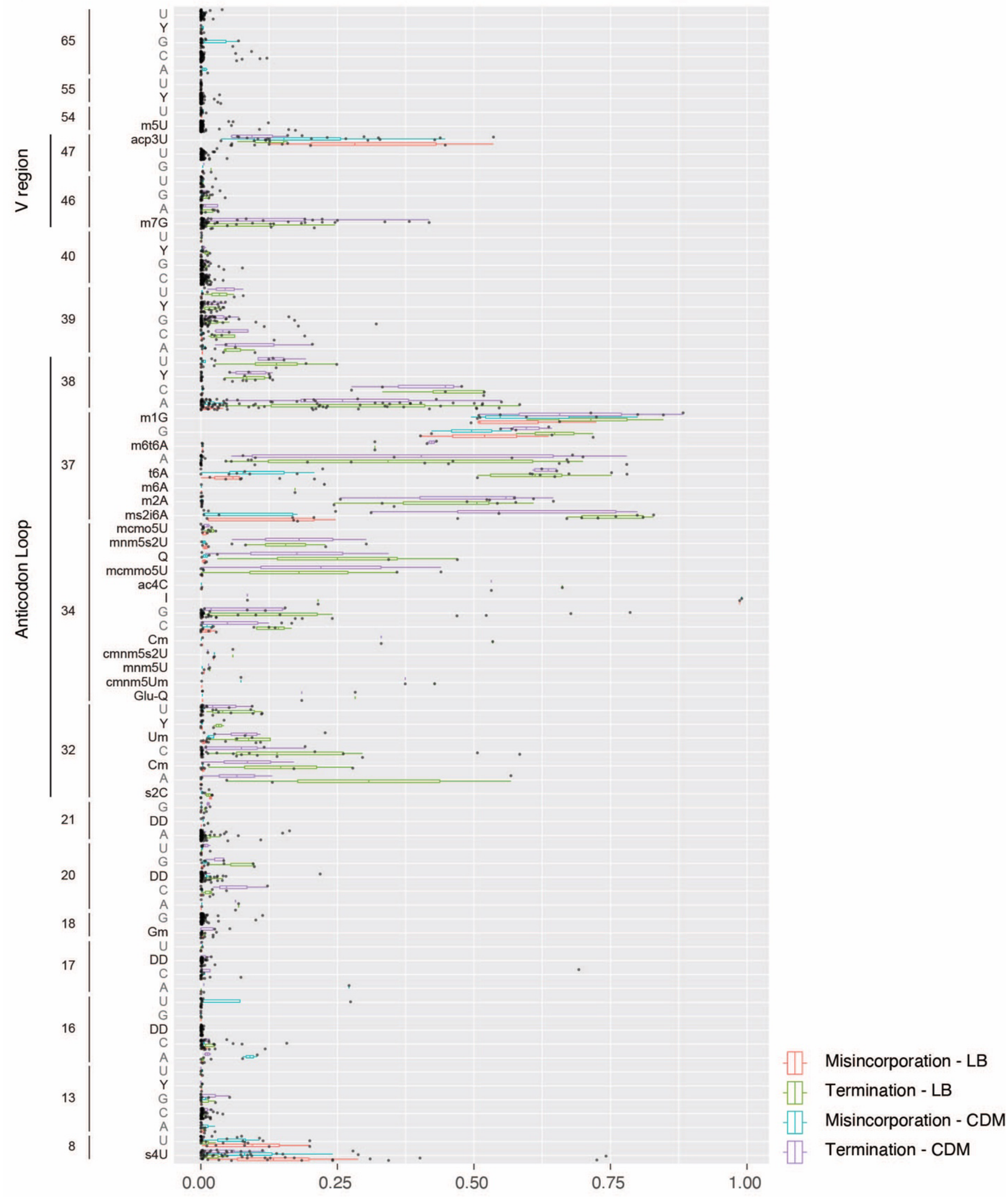


**Supplemental Figure 3.**

**(A)** Box and whiskers representation of average frequency of misincorporation and average termination score for each nucleotide found at positions annotated to include at least one modification. Positions included have a read coverage above 100 in both LB and chemically defined media (CDM) samples. Short name for each modification is used (Supplemental Table 11).

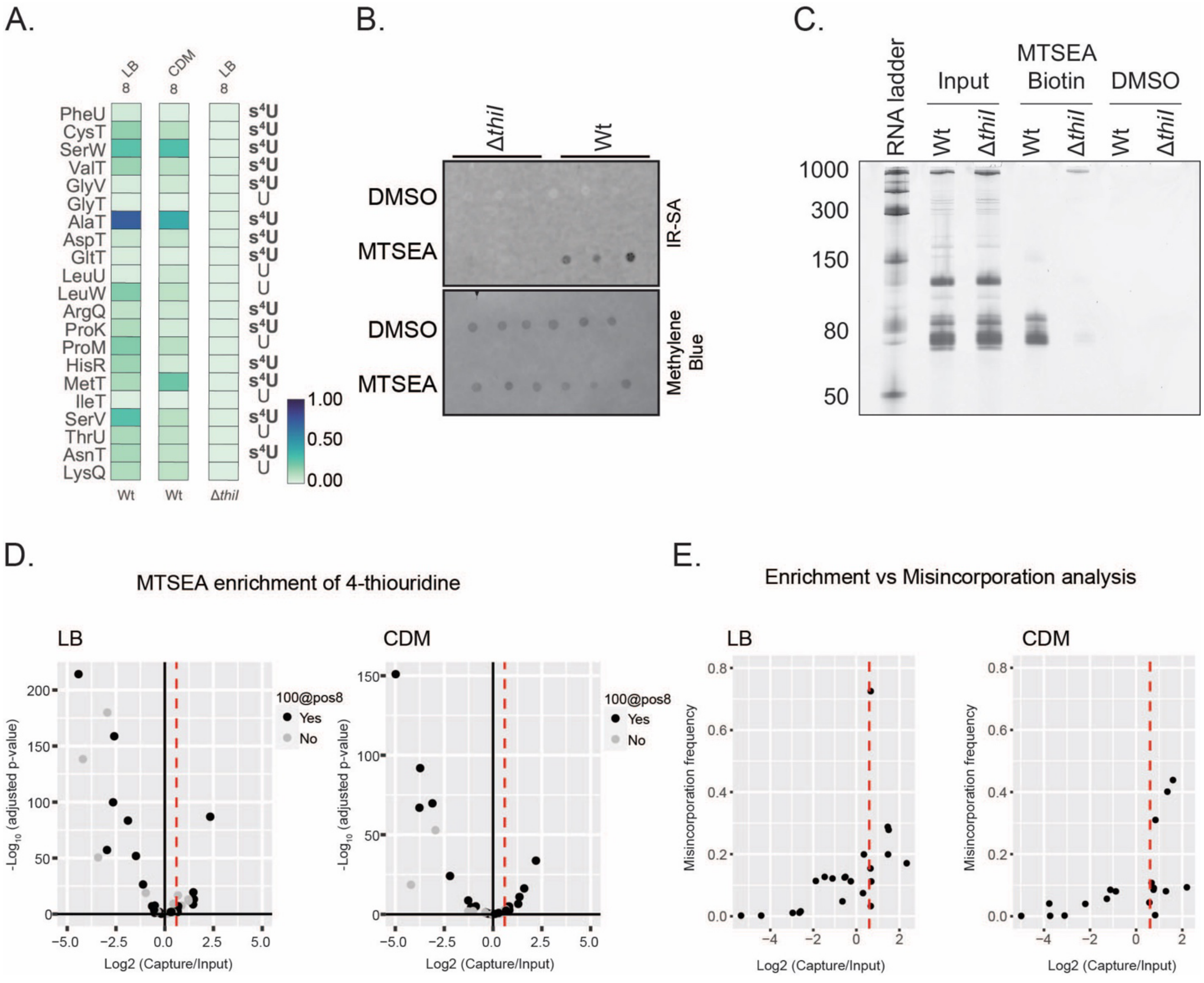


**Supplemental Figure 4.**

**(A)** Heatmap of average frequency (from three independent samples) of misincorporation at position 8 of tRNAs in *E. coli* wild type cells cultured in LB (left) or chemically defined media (CDM) (center) and Δ*thiI* mutant cells cultured in LB media (right). All cells were collected at $(OD)_{600}$=0.6. The color key at the right represents the frequency of misincorporation. Only tRNAs with read coverage > 100 at position 8 in all 3 experiments are represented. **(B)** Streptavidin (SA) dot blot was performed on RNA extracted from wild type (BW25113 strain) or Δ*thiI E. coli* cells and treated with DMSO or MTSEA. Methylene blue was used as a loading control. **(C)** RNA extracted from wild type and Δ*thiI E. coli* cells resolved on a 10% denaturing urea polyacrylamide gel. RNA was treated with MTSEA or DMSO, incubated with SA beads, and eluted RNA resolved in the gel.

Total RNA from each bacterial strain is included. **(D)** Volcano plot representation of log 2-fold change in abundance after MTSEA capture of tRNA extracted from *E. coli* cells cultured in LB (left) or chemically defined media (right). Dotted red line indicates the threshold used to define a tRNA as enriched. Color indicates if read coverage at position 8 is above 100. **(E)** Scatter plot representation of log 2-fold change in abundance after MTSEA capture and misincorporation frequency of tRNA from *E. coli* cells cultured in LB (left) or chemically defined media (right). Dotted red line indicates the threshold used to define a tRNA as enriched.

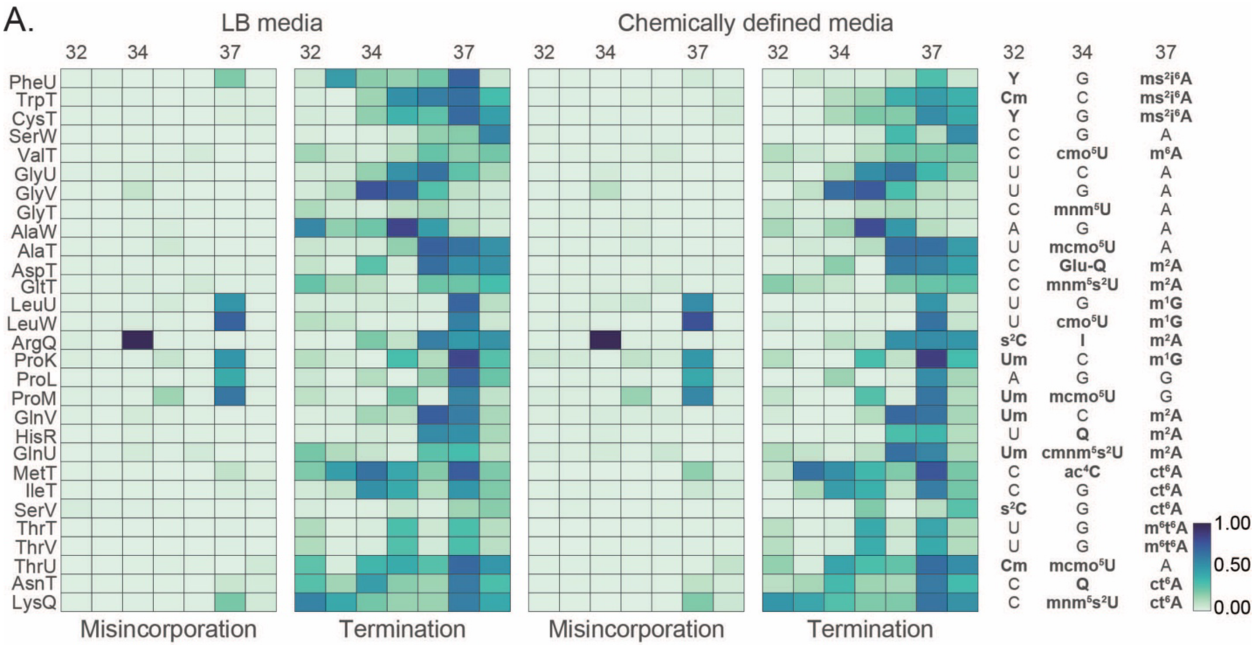


**Supplemental Figure 5.**

**(A)** Heatmap representation of average (from three independent samples) frequency of misincorporation or termination score between positions 32 and 38 in tRNA from *E. coli* cultured in LB (left) or chemically defined media (right). Only tRNAs with read coverage > 100 in the anticodon loop in both experiments are represented. Positions 32, 34 and 37, known to be modified are annotated at the top of each heatmap. Known modifications for E. coli are represented on the right. Short name for each modification is used (Supplemental Table 11). The color key at the right represents the frequency of misincorporation or termination score.

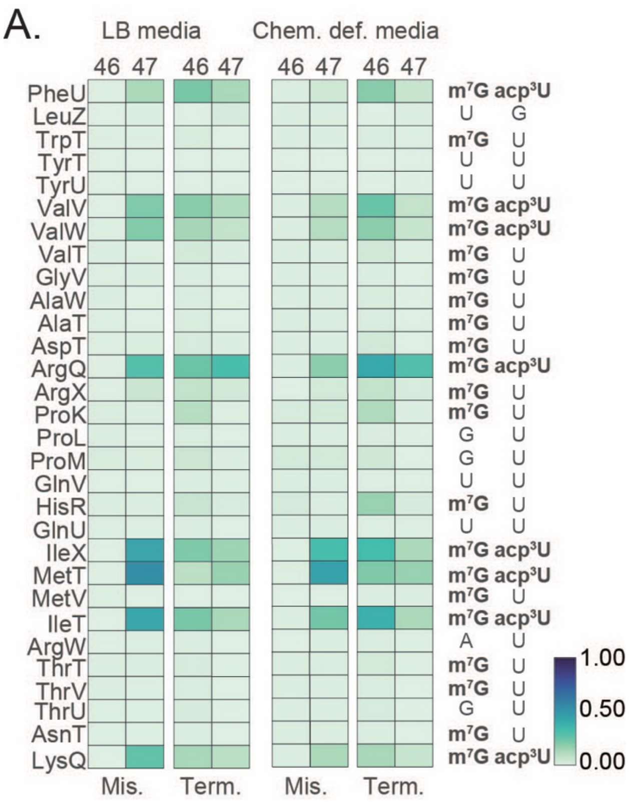


**Supplemental Figure 6.**

**(A)** Heatmap representation of average (from three independent samples) frequency of misincorporation and termination score at positions 46 and 47 in tRNA from *E. coli* cultured in LB (left) or chemically modified media (right) and collected at $(OD)_{600}$=0.6. Only tRNAs with read coverage > 100 at position 46 in both experiments are represented. Annotated nucleotide is represented to right side of the heatmap. Short name for each modification is used (Supplemental Table 11). The color key at the right represents the frequency of misincorporation or termination score.

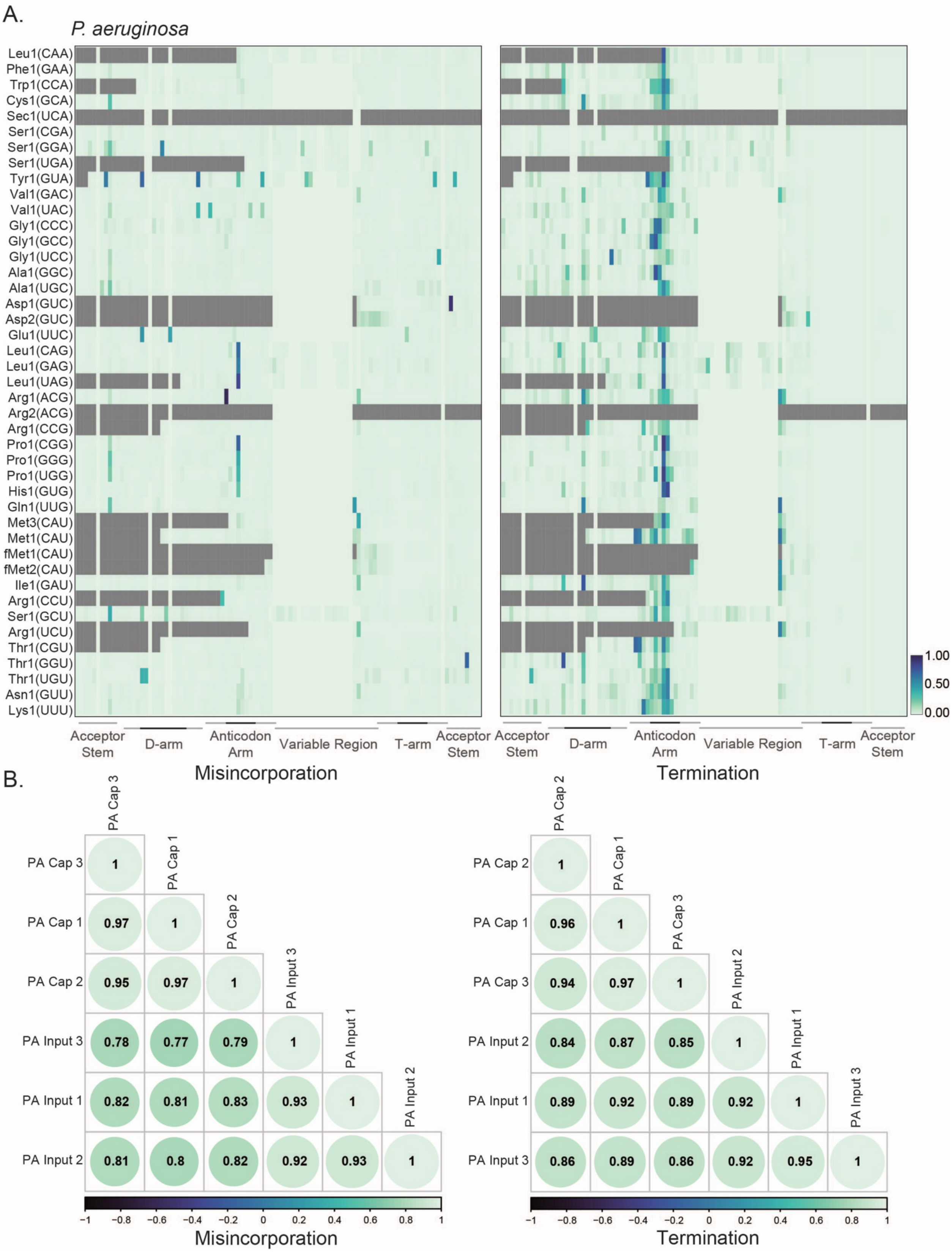


**Supplemental Figure 7.**

**(A)** Heatmap representation of average (from three independent samples) frequency of misincorporation (left) or termination score (right) at each position for *P. aeruginosa* tRNAs. All tRNAs in the custom transcriptome are represented. RNA was collected from cells cultured in chemically modified media and collected at $(OD)_{600}$=0.6. The color key at the right represents the frequency of misincorporation or termination score. Different regions of the tRNA are represented at the bottom. Darker lines indicate the loop in that region. The GtRNAdb names are used for *P. aeruginosa* tRNAs. The anticodon is shown in superscript. **(B)** Correlation plot for misincorporation and termination between replicates for total RNA and MTSEA-captured samples in *P. aeruginosa*.

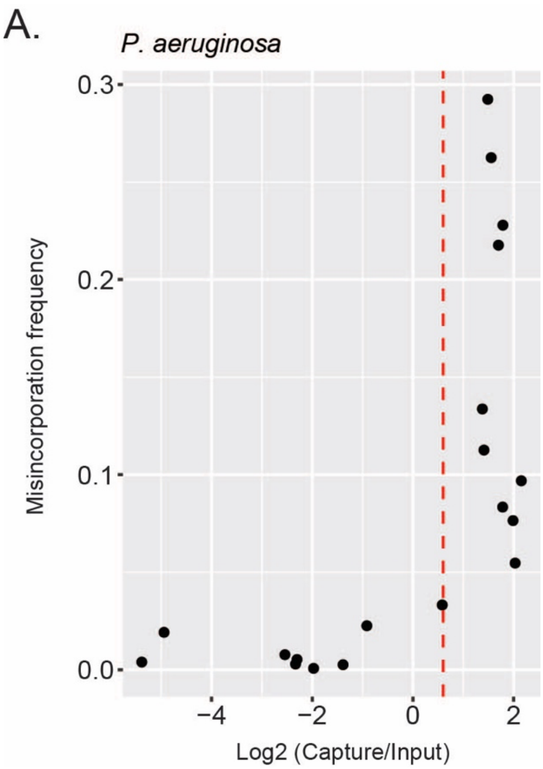


**Supplemental Figure 8.**

**(A)** Scatter plot representation of log 2-fold change in abundance after MTSEA capture and misincorporation frequency of tRNA from *P. aeruginosa* cells cultured in chemically defined media. Dotted red line indicates the threshold used for enrichment in the MTSEA enrichment experiment.

A.

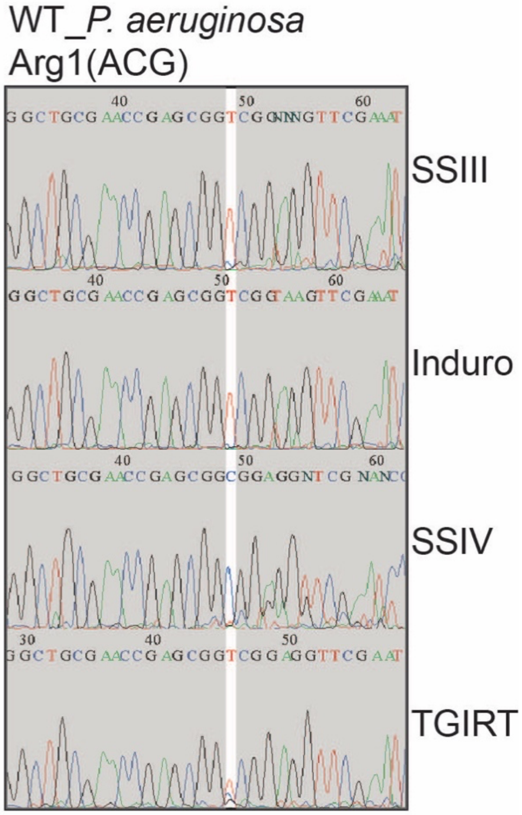


**Supplemental Figure 9.**

**(A)** Sanger sequencing chromatograms of targeted RT-PCR to detect the acp$^3$U signature observed at position 47 in tRNA$^{Arg1(ACG)}$ from *P. aeruginosa*. Listed are four reverse transcriptase enzymes that were tested.

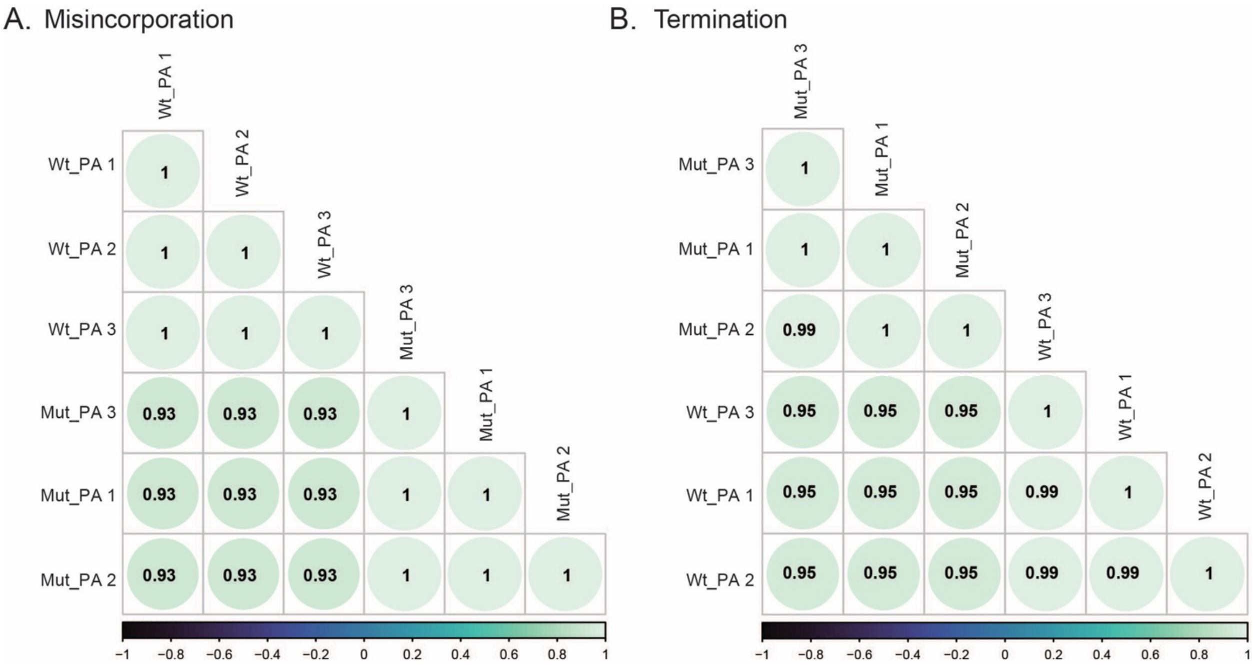


**Supplemental Figure 10.**

**(A, B)** Correlation plot for misincorporation and termination between replicates for wild-type (Wt_PA) and PA14_17650 mutant (Mut_PA) from *P. aeruginosa* RNA sequencing.

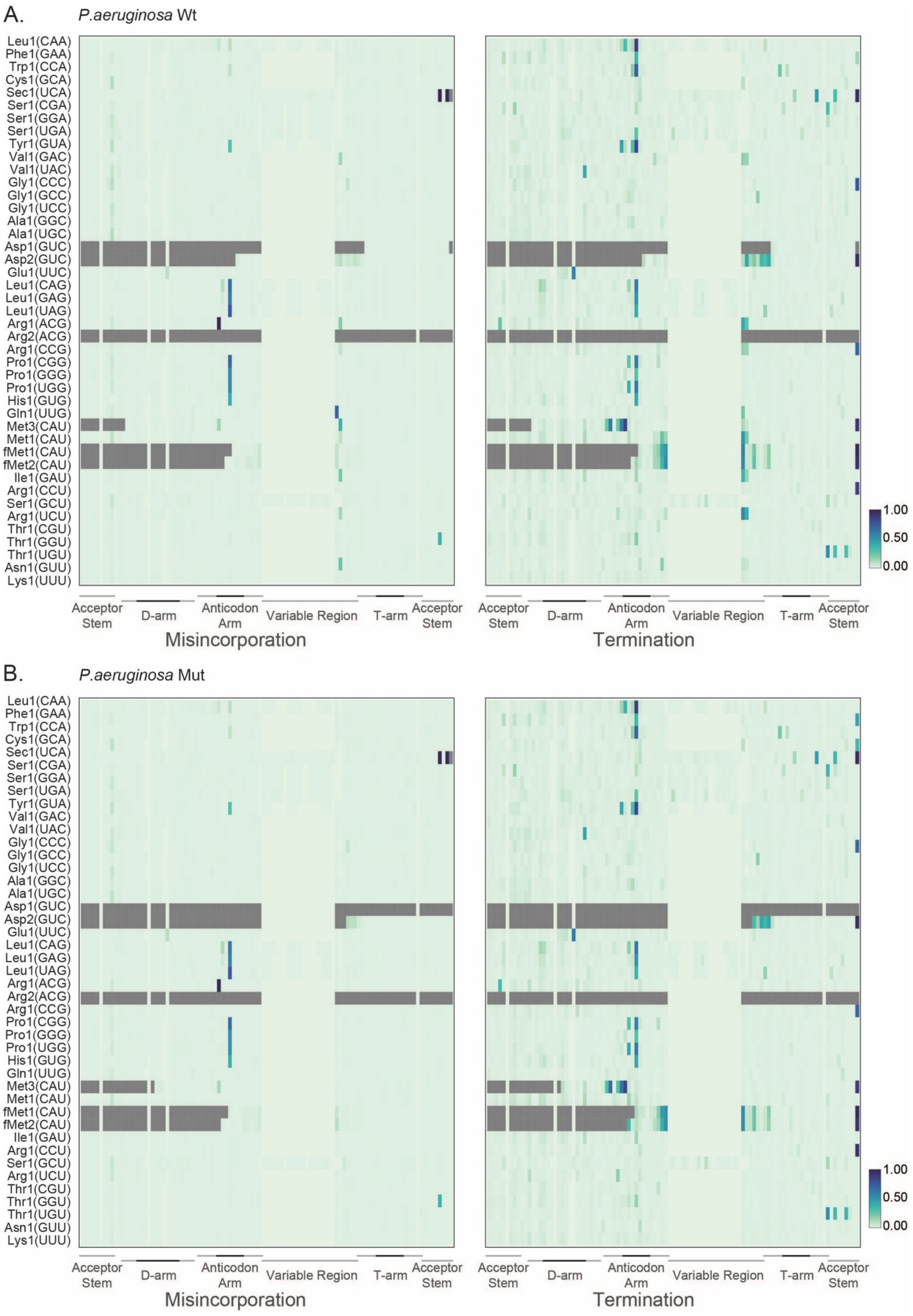
A.
P.aeruginosa Wt
Leu1(CAA)
Phe1(GAA)
Trp1(CCA)
Cys1(GCA)
Sec1(UCA)
Ser1(CGA)
Ser1(GGA)
Ser1(UGA)
Tyr1(GUA)
Val1(GAC)
Val1(UAC)
Gly1(CCC)
Gly1(GCC)
Gly1(UCC)
Ala1(GGC)
Ala1(UGC)
Asp1(GUC)
Asp2(GUC)
Glu1(UUC)
Leu1(CAG)
Leu1(GAG)
Leu1(UAG)
Arg1(ACG)
Arg2(ACG)
Arg1(CCG)
Pro1(CGG)
Pro1(GGG)
Pro1(UGG)
His1(GUG)
Gln1(UUG)
Met3(CAU)
Met1(CAU)
fMet1(CAU)
fMet2(CAU)
Ile1(GAU)
Arg1(CCU)
Ser1(GCU)
Arg1(UCU)
Thr1(CGU)
Thr1(GGU)
Thr1(UGU)
Asn1(GUU)
Lys1(UUU)
1.00
0.50
0.00
Acceptor Stem
D-arm
Anticodon Arm
Variable Region
T-arm
Acceptor Stem
Misincorporation
Termination
B.
P.aeruginosa Mut
Leu1(CAA)
Phe1(GAA)
Trp1(CCA)
Cys1(GCA)
Sec1(UCA)
Ser1(CGA)
Ser1(GGA)
Ser1(UGA)
Tyr1(GUA)
Val1(GAC)
Val1(UAC)
Gly1(CCC)
Gly1(GCC)
Gly1(UCC)
Ala1(GGC)
Ala1(UGC)
Asp1(GUC)
Asp2(GUC)
Glu1(UUC)
Leu1(CAG)
Leu1(GAG)
Leu1(UAG)
Arg1(ACG)
Arg2(ACG)
Arg1(CCG)
Pro1(CGG)
Pro1(GGG)
Pro1(UGG)
His1(GUG)
Gln1(UUG)
Met3(CAU)
Met1(CAU)
fMet1(CAU)
fMet2(CAU)
Ile1(GAU)
Arg1(CCU)
Ser1(GCU)
Arg1(UCU)
Thr1(CGU)
Thr1(GGU)
Thr1(UGU)
Asn1(GUU)
Lys1(UUU)
1.00
0.50
0.00
Acceptor Stem
D-arm
Anticodon Arm
Variable Region
T-arm
Acceptor Stem
Misincorporation
Termination

**Supplemental Figure 11.**

**(A)** Heatmap of average frequency (from three independent samples) of misincorporation (left) or termination score (right) at each position for wild-type (Wt) *P.aeruginosa* and *P.aeruginosa* PA14_17650 (Mut) mutant tRNAs extracted from cells cultured in chemically modified media (CDM) and collected at $(OD)_{600}$=0.6. All *P.aeruginosa* tRNAs are represented with the GtRNAdb names, and the anticodons shown in parentheses. The color key at the right represents the frequency of misincorporation or termination score. The grey refers to regions of less than 100 reads per position. Different regions of the tRNA are represented at the bottom from the 5′ (left) to the 3′ end (right) of the tRNA. Darker lines indicate the loop in that region. **(B)** Heatmap of average frequency (from three independent samples) of misincorporation (left) or termination score (right) at each position for PA14_17650 mutant (Mut) *P. aeruginosa* tRNAs extracted from cells cultured in LB and collected at $(OD)_{600}$=0.6. All *P.aeruginosa* tRNAs are represented with the GtRNAdb names, and the anticodons shown in parentheses. The color key at the right represents the frequency of misincorporation or termination score. The grey refers to regions of less than 100 reads per position. Different regions of the tRNA are represented at the bottom from the 5′ (left) to the 3′ end (right) of the tRNA. Darker lines indicate the loop in that region.

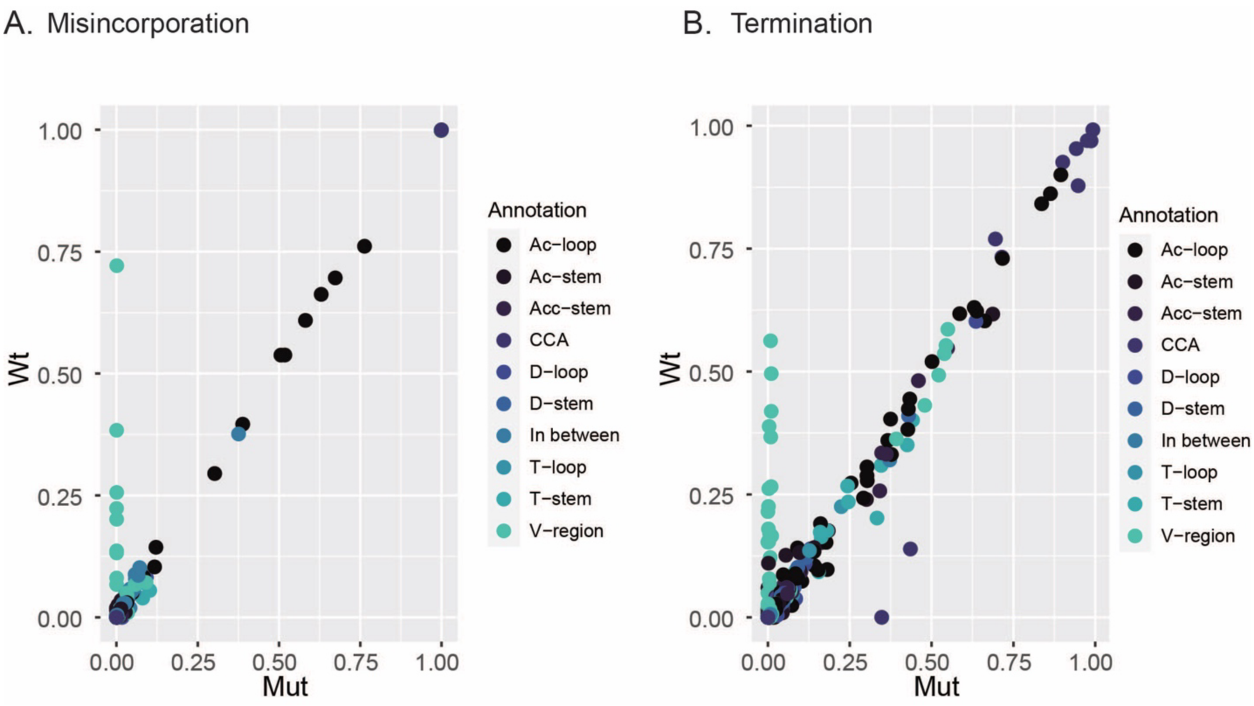


**Supplemental Figure 12.**

**(A, B)** Scatter plots representing the levels of misincorporation or termination scores that compare at all positions between wild-type (Wt) or PA14_17650 mutant (Mut) tRNA. Color coding at the right of each graph indicates the location of the tRNA positions within the annotated region of the tRNA.

**Supplemental Table legends:**

**Supplemental Table 1.** *E. coli* tRNAs are listed, including transcript ID, tRNA name, anticodon and tRNA sequences. tRNAs that contained 100% sequence identity were grouped together and named by the first tRNA in the group.

**Supplemental Table 2.** Master table of *E. coli* tRNA misincorporation and termination scores for each position from all tRNA of cultures grown in LB and CDM. Listed are the transcript ID, tRNA name, annotated position and corresponding region (a zero refers to locations in which regional information is not available), the reference nucleotide at each position, all annotated modifications, average coverage depth, and misincorporation (mis) and termination (ts) scores according to our sequencing data. Scores were calculated only for depths greater than 100.

**Supplemental Table 3.** Deseq analysis output of MTSEA-biotin captures versus inputs from E. coli grown in LB (3A) or CDM (3B). Listed are the transcript ID, tRNA name, base mean, log2fold change, lfc standard error value, the Wals statistic (stat), p values and adjusted p values (padj).

**Supplemental Table 4.** Table of E. coli tRNA misincorporation and termination scores for positions 32, 34 and 37 from all tRNA of cultures grown in LB and CDM. Listed are the transcript ID, tRNA name, annotated position and corresponding region, the reference nucleotide at each position, all annotated modifications, average coverage depths, and misincorporation (mis) and termination (ts) scores according to our sequencing data. Scores were calculated only for depths greater than 100.

**Supplemental Table 5.** Table of E. coli tRNA misincorporation and termination scores for positions 46 and 47 from all tRNA of cultures grown in LB and CDM. Listed are the transcript ID, tRNA name, annotated position and corresponding region, the reference nucleotide at each position, all annotated modifications (a zero refers to no annotated nucleotide at that position), average coverage depths, and misincorporation (mis) and termination (ts) scores according to our sequencing data. Scores were calculated only for depths greater than 100.

**Supplemental Table 6.** P. aeruginosa tRNA gene information pulled from Ensembl (6A). tRNAs transcript ID, P. aeruginosa (PA) gene name, and assigned tRNA nomenclature for each tRNA (6B). Conserved E. coli and P. aeruginosa tRNAs were matched based on anticodons and by sequence similarity (6C).

**Supplemental Table 7.** Master table of matched *E. coli* and *P. aeruginosa* tRNAs. Listed are the transcript ID, tRNA name, annotated position and corresponding region (a zero refers to locations in which regional information is not available), the reference nucleotide at each position, all annotated modifications, average coverage depths, and misincorporation (mis) and termination (ts) scores according to our sequencing data. Scores were calculated only for depths greater than 100.

**Supplemental Table 8.** Deseq analysis output of MTSEA-biotin captures versus inputs of P. aeruginosa. Listed are the transcript ID, tRNA name, base mean, log2fold change, lfc standard error value, the Wals statistic (stat), p values and adjusted p values (padj).

**Supplemental Table 9.** Data used for the plot in Supplemental Figure 8 indicating a positive correlation between misincorporation score and log 2-fold enrichment by MTSEA-biotin. Listed are the transcript ID, tRNA name, misincorporation scores (mis) and log2fold values (l2f) for each tRNA that had 100 reads or more at position 8 and significant padj values ($< 0.05$).

**Supplemental Table 10.** Data from the correlation plot in Figure 2C comparing log2fold enrichment values between each matched tRNA of *E. coli* (Ec) and *P. aeruginosa* (PA) MTSEA-biotin capture experiments. Listed are each anticodon and the subsequent log2fold values.

**Supplemental Table 11.** Listed are the all the different modifications found in tRNA and their corresponding abbreviations or short names, based on literature and the Modomics RNA modification database [8,36].

**Supplemental Table 12.** List of tRNA modifying enzymes from *E. coli* and *P. aeruginosa* and their homology BLAST scores.

**Supplemental Table 13.** Master table of *P. aeruginosa* tRNAs from wild-type (Wt) and PA14_17650 (*tapT*) mutant (Mut) sequencing data. Listed are the transcript ID, tRNA name, reference base, annotated position and corresponding region (a zero refers to locations in which regional information is not available), the anticodon (AC), and the average misincorporation (mis), depths, and termination scores according to the sequencing data. Scores were calculated only for depths greater than 100, otherwise listed as NA.

**Supplemental Table 14.** Oligo sequences used to make sequencing libraries and for targeted RT-PCR and sanger sequencing.
